# A single-cell transcriptomic atlas of peripheral blood immune cells spanning progressive canine leishmaniosis

**DOI:** 10.64898/2025.12.18.695149

**Authors:** Danielle P. Uhl, Daniel J. Holbrook, Max C. Waugh, Shoumit Dey, Najmeeyah Brown, Karen I. Cyndari, Jacob J. Oleson, Paul M. Kaye, Christine A. Petersen

**Affiliations:** Department of Epidemiology and Center for Emerging Infectious Diseases, College of Public Health, University of Iowa, Iowa City, IA, USA; Department of Obstetrics and Gynecology, College of Medicine, University of Iowa, Iowa City, IA, USA; Hull York Medical School and York Biomedical Research Institute, University of York, York, UK; College of Veterinary Medicine, The Ohio State University, Columbus, OH, USA; Department of Emergency Medicine, University of Iowa Hospitals and Clinics, Iowa City, IA, USA; Department of Biostatistics, College of Public Health, University of Iowa, Iowa City, IA, USA

## Abstract

Dogs play a major role in sustaining transmission of *Leishmania infantum* to people, thus prevention and treatment of canine leishmaniosis (CanL) to reduce transmission represents an unmet public health need. Although advances have been made in understanding how immunopathology correlates to infectiousness as CanL progresses, the immune mechanisms underlying transition between disease stages and terminal decline remain ill-defined. To address this knowledge gap, we generated a comprehensive atlas of peripheral immune cells from control dogs and naturally-infected dogs using single-cell RNA sequencing. The LeishDog Atlas captures the cellular and transcriptional complexity underlying CanL, tracing shifts in immune composition and gene expression across progressive, well-defined disease stages. Notably, we identified distinct myeloid, CD4^+^ and CD8^+^ T cell phenotypes and transcriptional states associated with disease progression. This resource provides a valuable framework for understanding systemic immune dysregulation in CanL, including determinants of T cell exhaustion and myeloid cell activation, and establishes a foundation for comparative, translational, and mechanistic studies of CanL immunopathology and transmissibility.

## INTRODUCTION

Human visceral leishmaniasis (VL; kala azar) is a vector-borne neglected tropical disease caused by the obligate intracellular protozoans *Leishmania infantum* and *L. donovani*. A total of 79 countries were considered endemic in 2023, with an estimated 50,000 – 90,000 new cases and 20,000 reported deaths annually^1–3^. Following decades of elimination campaigns on the Indian Subcontinent, the greatest burdens of VL are now found primarily in East Africa and Brazil, with cases also occurring throughout the Mediterranean Basin^2^. Socioeconomic determinants such as poverty, malnutrition, poor housing, and limited access to healthcare and public health services critically influence both susceptibility to parasite infection and sand fly exposure^2^. Currently, there is no vaccine for preventing human disease and available chemotherapeutic options are limited and hindered by issues of toxicity, emerging resistance, and incomplete parasite clearance^4–6^. Transmission of *L. donovani* is largely regarded as anthroponotic, though recent evidence supports a canine reservoir^7^. In contrast, transmission of *L. infantum* in Brazil and the Mediterranean basin is zoonotic^8,9^.

For zoonotic VL, domestic dogs are critical hosts that sustain transmission to humans through sand fly vectors^10–12^. Dogs are also considered effective sentinel hosts, with disease prevalence closely mirroring human exposure in endemic areas^13^. Although dogs with advanced clinical signs and increased parasitemia were assumed to be the most infectious to sand flies^14,15^, several studies demonstrate that dogs with mild canine leishmaniosis (CanL) can be more infectious to sand flies^16,17^. *L. infantum* infection can also be maintained through vertical transmission^18,19^. However, regardless of route of transmission, common clinical manifestations of CanL include lethargy, weight loss, anemia, skin lesions, lymphadenopathy, and hepatosplenomegaly. As in human VL, CanL is lethal if left untreated, due in part to immune complex-mediated renal failure^20^.

Protective immunity to *L. infantum* infection in humans and dogs depends on a balance between inflammatory and regulatory immune responses^21,22^. Pro-inflammatory type one (T_H_1) responses, mainly sustained by IFNγ-producing CD4^+^ T cells, are critical for activating microbicidal mechanisms that restrain parasite replication^22,23^. Cytotoxic CD8^+^ T cells may assist in controlling infection by killing infected host cells or contributing to the proinflammatory cytokine milieu^24–26^. In contrast, *Leishmania*-specific antibodies may facilitate phagocytosis of parasites, while non-specific IgG antibodies, notably in CanL, contribute directly to end-stage disease^27,28^. Other suppressive mechanisms including type 1 regulatory (T_R_1) CD4^+^ T cells and regulatory B cells also emerge during advanced clinical disease that further dampen host microbicidal functions through IL-10 production^22,29–32^. Numerous studies have used transcriptomics of whole blood or PBMCs to expand our understanding of the immune status of patients with active VL and after treatment^33,34^. These confirm systemic alterations in immune cell frequencies, cytokine production, and host cell metabolism, and have identified novel pathways associated with CD4^+^ and CD8^+^ T cell activation and exhaustion^35–37^. However, these high-resolution approaches have not yet been applied to better understand host immunity during early CanL disease or factors that may relate to disease progression.

Across hosts, very little is known about the immune mechanisms underlying transition from asymptomatic infection to clinical disease and subsequently leading to terminal decline. Therefore, we aimed to define peripheral immune cell composition and transcriptional programs associated with disease progression across defined clinical stages of CanL using single-cell RNA sequencing (scRNA-seq). Here, we describe the LeishDog Atlas, the first single-cell transcriptomic atlas of peripheral blood immune cells from dogs naturally-infected with *L. infantum.* The Atlas provides a comprehensive view of the canine peripheral blood immune system and spans the distinct clinical stages of CanL defined by LeishVet guidelines (LV1-LV4)^20^. Our data demonstrates for the first time the dynamics and transcriptional heterogeneity within the circulating lymphoid and myeloid cell populations in CanL, and identifies transcriptional changes associated with disease progression. The LeishDog Atlas provides a high-resolution overview of systemic immune alterations during CanL progression that will inform fundamental mechanistic studies and help future efforts towards developing transmission-preventing interventions, vaccines, and immunotherapies. In addition, it represents a significant resource for those working in related fields of canine immunology and infection.

## RESULTS

### Study design and analysis of immune cells

To characterize immune cells across the spectrum of CanL, we performed scRNA-seq on PBMCs from dogs naturally infected with *L. infantum* by vertical transmission and from uninfected dogs (NLC) (**Figure 1A; Table S1)**. A total of 68,323 cells from 16 PBMC samples passed QC assessment (NLC, 8,641 cells; LV1, 10,882 cells; LV2, 18,914 cells; LV3, 13,111 cells; LV4, 16,775 cells). After normalization and integration, unsupervised clustering (Level 1 clustering) identified 15 cell clusters projected into a uniform manifold approximation and projection (UMAP) plot (**Figure 1B**). Canonical marker expression was used to identify T cells (*CD3E*, *TRAV9-2* (LOC607937), and *TRBC1* (LOC480788)); Natural Killer (NK) cells (*KLRD1*, *KLRK1*, and *NCR3*); B cells (*CD79A*, *MS4A1*, and *CD19*); monocytes (*LYZ*, *CSF1R*, and *CTSS*); dendritic cells (DCs; *CD1D*, *PLD4*, and *FLT3*); neutrophils (*PADI4*, *CSF3R*, and *MEGF9*); eosinophils (*IL5RA*, *CCR3,* and *ITGAM*); basophils (*IL3RA*, *MS4A2*, and *CPA3*); platelets (*PPBP*, *TUBB1*, and *GP93*); and plasma cells (*JCHAIN, MZB1, IRF4*) (**Figure 1C-D**). Clustering was used to assign the cell types to three major groups: T/NK cells (*n* = 42,011 cells), B cells (*n* = 8,275 cells), and myeloid cells (*n* = 18,037 cells) (**Figure 1E**). Low-resolution clustering initially grouped plasma cells with T/NK cells, therefore manual re-assignment as B cells was performed (see **STAR**⍰**METHODS**). The identification of low-density neutrophils and other granulocytes in our integrated dataset was anticipated^38–40^. To further support our global cell type annotation, we used reference mapping against the pre-annotated human PBMCs dataset from the Azimuth database^41^ (see **STAR**⍰**METHODS)**. Human reference mapping aligned substantially with our annotation for canine mononuclear cell populations (**Figure 1F**).

**Figure 1.**
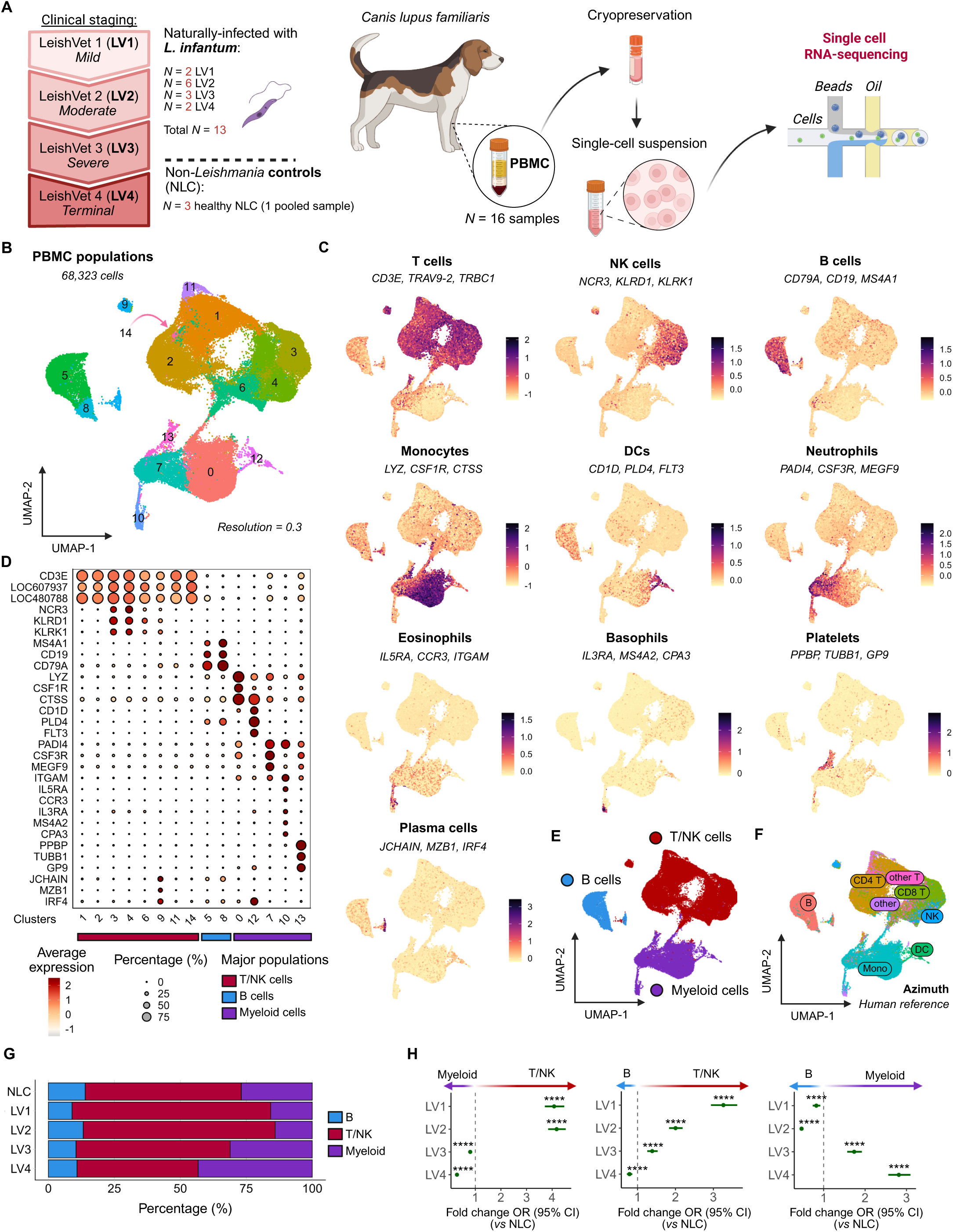
Immune cell annotation and T/NK-myeloid lineage dynamics in CanL PBMC single-cell transcriptomes. **(A)** Schematic overview of the study design showing peripheral blood mononuclear cell (PBMC) collection from dogs and the subsequent single-cell RNA-seq workflow. Created in BioRender. Uhl, D. (2025) https://BioRender.com/ktukw9j. **(B)** UMAP visualization showing 15 clusters of canine peripheral blood immune cells. **(C)** Feature plots showing module scores for selected canonical marker genes used to identify and distinguish immune cell populations. **(D)** Dot plot showing cluster expression and proportion of canonical marker genes (LOC607937: *TRAV9*-1; LOC480788: *TRBC1*). **(E)** UMAP showing manual annotation of three major PBMC populations (Level 1). **(F)** UMAP displaying predicted labels for major PBMC populations mapped using the Azimuth human PBMC reference dataset. **(G)** Bar plot showing the distribution of major immune populations across CanL clinical stages, colored by population. See also Table S2. **(H)** Forest plots of the fold-change in odds ratio (OR) of each major population abundance across clinical stages relative to NLC. Points indicate estimated ORs and whiskers represent 95% confidence intervals from a mixed-effects logistic regression model with random intercepts for individual dogs. P values were adjusted for multiple comparisons using the Benjamini–Hochberg method (****adjusted *P* < 0.0001). See also Table S3.

We compared the distribution of T/NK, B, and myeloid cells for NLC dogs and for CanL dogs from LV1 (mild) to LV4 (terminal) disease stage (**Figure 1G, Table S2**). Using a mixed-effects logistic regression approach with random intercept per sample (model details in **STAR**⍰**METHODS**), aligned with prior works^42,43^, we confirmed that when compared to NLC, T/NK cells predominated in early stages of disease whereas myeloid cells became more prominent in advanced disease (**Figure 1H, Table S3)**. In contrast, overall B cell frequencies within peripheral blood did not change significantly across LV stage (**Figure 1G**). These findings highlight a dynamic reorganization of the peripheral immune compartment over the course of CanL progression. To better understand the mechanisms that might underlie these changes, the three *Level 1* immune populations were then independently re-clustered (*Level 2*) (**Figure S1A**).

### B cells

In CanL, B cells generate both *Leishmania-*specific and polyclonal antibodies, with varying host protective and disease promoting roles^22^. A total of nine distinct B cell subpopulations were identified by unsupervised clustering (**Figure 2A** and **Table S4**). Four clusters were naïve B cells (*n* = 4,368 cells; *PAX5, BACH2*). As these were differentiated mainly by variable usage of immunoglobulin light chain genes (**Figure S3A and Table S4**), these clusters were pooled as a single naïve B cell population. Other B cell clusters were identified as transitional B cells (TrB; *n* = 410 cells; *VPREB3*, *IGFR1*, *SOX4, IGHM* (LOC100685971)*, CD79A/B*)), unswitched B cells (*n* = 841 cells; *CD44, IGHM, BACH2*), two populations of class-switched B cells (B_1, *n* = 1,210; B_2, *n* = 484 cells), both lacking *IGHM* but having numerous differentially expressed genes (**Figure S3B-D and Table S4**), and terminally differentiated plasma cells (PC; *n* = 258 cells; *PRDM1*, *IRF4*, *JCHAIN, MZB1*) (**Figure 2B-C**). Memory B cells could not be formally identified due to the lack of significant *CD27* expression across clusters (**Figure S3E**). Two clusters (#7, *n* = 402; #9, *n* = 302) were excluded from further analysis, being subsequently identified as doublets (**Figure S2**; see **STAR**⍰**METHODS**). As with total B cell subpopulations, we observed minimal differences in the frequency of these defined B cell populations across disease stage (**Figure 2D and Table S2**).

**Figure 2.**
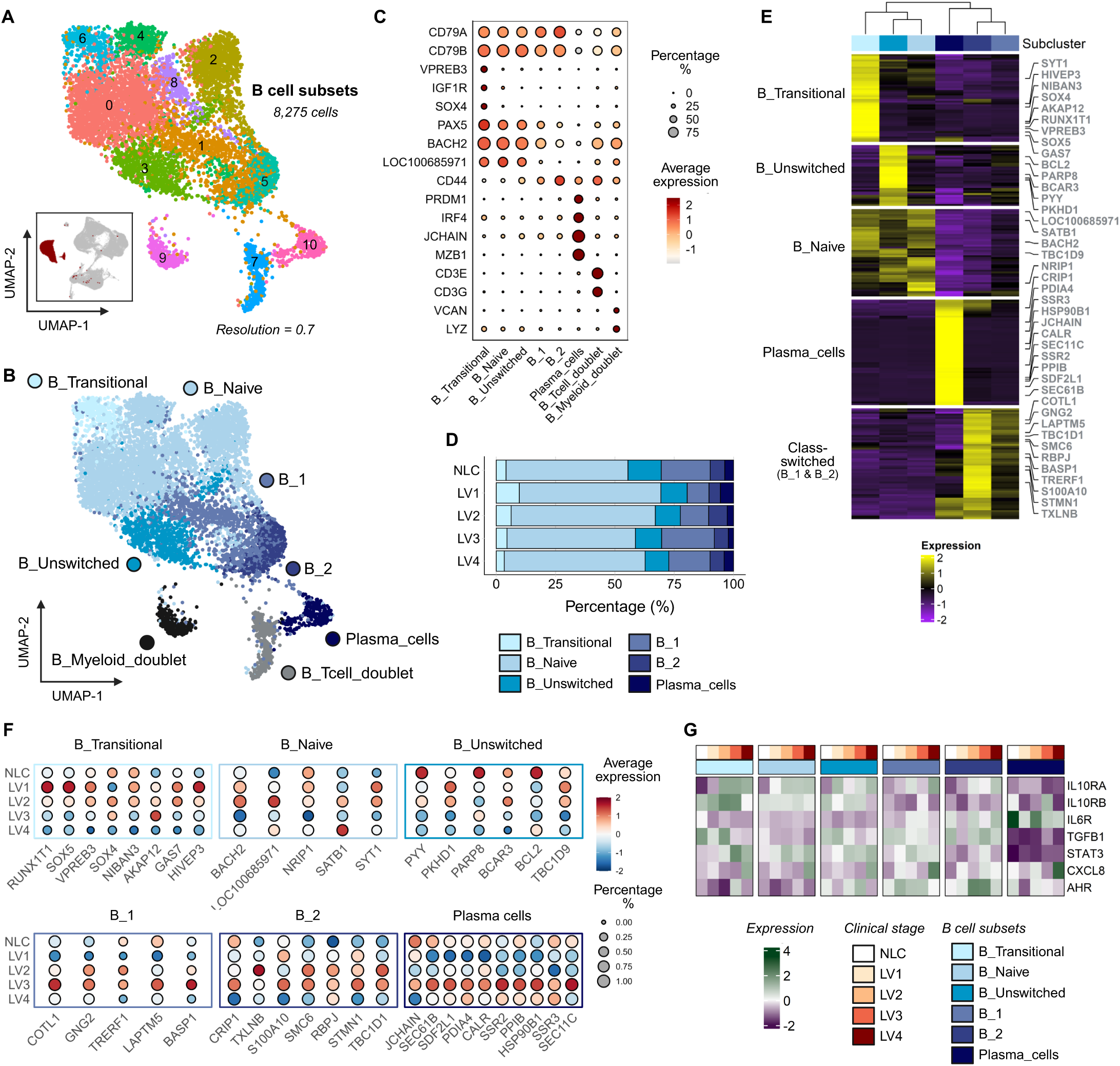
Annotated B cell subpopulations in CanL exhibit minimal frequency change but marked transcriptional shifts during progression. **(A)** UMAP visualization of 11 B cell subclusters. The inset shows the projection of these cells onto the PBMC UMAP. **(B)** UMAP visualization of 8 annotated populations of B cells. **(C)** Dot plot showing canonical marker gene expression and proportions in B cell subclusters (LOC100685971: *IGHM*). **(D)** Bar plot depicting the proportion of cells across CanL clinical stages, colored by B cell population. See also Table S2. **(E)** Heatmap visualization of top 50 differentially expressed genes (DEGs) of each annotated B cell subcluster. DEGs and subclusters arranged by hierarchical clustering. Highlighted DEGs correspond to the those visualized in Figure 2F. See also Table S4. **(F)** Dot plot showing expression and proportions within specific cell populations of curated gene sets across CanL clinical stages. **(G)** Heatmap of average expression for genes associated with inflammation and immunoregulation across B cell major subpopulations and CanL clinical stages.

Although frequencies varied little with disease progression, we noted that transcript abundance for many of the signature genes that define these populations varied across LV stage. For example, TrB cells at LV1 and LV2 had high transcript abundance for genes associated with immaturity and differentiation (*VPREB3* and *GAS7*) and stress-and inflammation-associated genes (*SOX5*, *HIVEP3*, and *NIBAN3*). Naïve B cells at LV2 had more abundant transcripts for *BACH2*, *SATB1* and *NRP1* (associated with maintenance of homeostasis and naïve B cell identity) and *SYT1* (associated with vesicle trafficking) when compared to naïve B cells in NLC and at later stages of disease. Transcripts for canonical PC genes, as well as B_1- and B_2-related transcripts, were most abundant at LV3 (**Figure 2E-F**).

B cells have been proposed to have regulatory properties during CanL and other infectious diseases^31,44^. Hence, we asked whether B cells exhibit transcriptional evidence of immune regulation or responsiveness to inflammatory signals during infection using a curated gene set representing these pathways (**Figure 2G**; see **STAR**⍰**METHODS**). Although TGF-β is a well-recognized immunosuppressive cytokine associated with regulatory B cells and VL^45,46^, transcript abundance declined across most B cell subsets as infection progressed, with the exception of B_2 cells, and was consistently low in both naïve B cells and PCs. IL-10 was not detected at the transcriptional level. However, *IL10RA* and its downstream target *STAT3* were notably more abundant at LV4 in all B cell populations except PCs, suggesting heightened responsiveness to IL-10. *IL6R* transcript abundance was low throughout disease. Transcript abundance for *CXCL8* (IL-8), a prominent neutrophil chemotactic cytokine, was greatest at LV4 in all B cell subsets except B_2 cells, suggesting that B cells may contribute to the inflammatory environment associated with terminal-stage disease. In contrast, transcripts for *AHR*, a transcription factor implicated in with immune regulation, metabolic adaptation, and suppression of effector responses in B cells^47^, were most abundant in unswitched B, B_1 and to a greater extent B_2 cells at earlier stages of disease progression. Together, these findings revealed that most peripheral B cell populations adopt overlapping immunoregulatory and inflammatory programs during CanL progression, with B_2 cells maintaining a distinct, non-inflammatory transcriptional profile.

### T cells and Innate lymphoid cells

T cells and innate lymphoid cells (ILCs) play pivotal roles in innate and cell-mediated immunity to *Leishmania*^21,22^. Level 2 clustering uncovered 35 subclusters (**Figure 3A**) that were broadly categorized as CD3^+^CD4^+^ (*CD4*; *n* = 17,903 cells), CD3^+^CD8^+^ (*CD8A*, *CD8B*; *n* = 16,188 cells), CD3^+^ double negative (DN; CD4^-^CD8^-^; *n* =3,399 cells), cytotoxic DN (*NKG7^+^; n* = 3,014) **(Figure 3B-C)**. Across the CanL spectrum, the proportions of CD4^+^ relative to CD8^+^, DN cells and cytotoxic DN T/NK cells declined significantly relative to NLC, while CD8^+^ cells predominated overall (**Figures 3D-E and Tables S2-S3**). These findings suggest a CD8□ T cell-biased immune profile emerging with disease progression. To further identify heterogeneity within these broadly defined T/NK cells, we performed further sub-clustering (Level 3; **Figure S1B**). A further subcluster identified as putative *Tcell_Myeloid_doublet* (*CD3*, *CD4, LYZ*, *VCAN*; *n* = 1,507) was excluded from further analysis (**Figure S2**; see **STAR**⍰**METHODS**).

**Figure 3.**
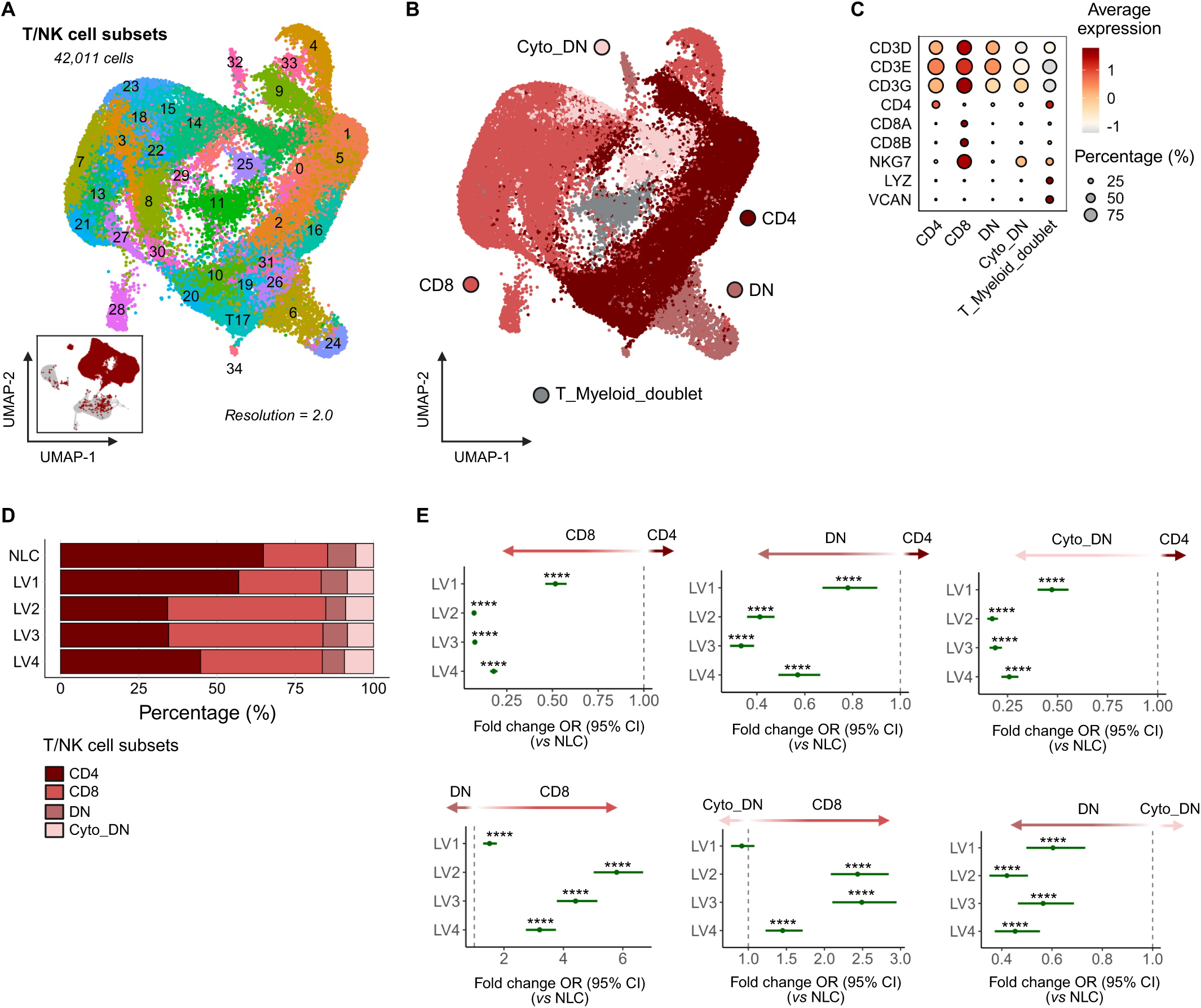
Annotation of T/NK cells (Level 2) revealed broad heterogeneity and a progressive shift toward CD8□ dominance over CD4^+^ T cells. **(A)** UMAP visualization of 35 T/NK cell subclusters. The inset shows the projection of these cells onto the PBMC UMAP. **(B)** Dot plot showing canonical marker expression and proportions of T/NK cell subclusters. **(C)** UMAP visualization showing manual annotation of 5 major T/NK cell populations (Level 2). **(D)** Bar plot displaying the proportion of CD4^+^, CD8^+^, DN, and cytotoxic DN T/NK cells across CanL clinical stages, colored by population. See also Table S2. **(E)** Forest plot of the fold-change in odds ratio (OR) of CD4^+^ over CD8^+^ T/NK cell abundance across clinical stages relative to NLC. Points indicate estimated ORs, and whiskers represent 95% confidence intervals from a mixed-effects logistic regression model with random intercepts for individual dogs. *P* values were adjusted for multiple comparisons using the Benjamini–Hochberg method (****adjusted *P* < 0.0001). See also Table S3.

### CD4^+^ and DN T cells

We identified 11 CD4^+^ and four DN T cells populations (**Figure 4A-D and Table S2**), namely naïve (CD4_Naive; *RSG10*, *CCR7*, *SELL*, *LEF1, TCF7*), central memory (CD4_T_CM_; *SELL*, *LEF1, TCF7, TSHZ2)*, effector memory (CD4_T_EM_; *CD28*, *LGALS1*, *ITGB1*), an intermediatory population (CD4_Int; lower expression of both T_CM_ and T_EM_markers), T_H_2-like T_EM_s (CD4_T_H_2; *CCDC3*, *SYTL3*, *LGALS3*, *ITGA2*), T_H_17-like T_EM_s (CD4_Th17; *RORA*, *ADAM12*, *CCR6*, *IL1R1*), T_H_1-like T_EM_s (CD4_Th1; *RCAN2*, *TBX21*, *IFNG*, *IL21*, *IL12RB2*), T_REG_s (CD4_T_REG_; *IKZF2*, *CTLA4*, *IL2RB*), a cytotoxic population (CD4_CTL; *CD4*, *IL7R*, *KLRG1*, *IL18R1*), and an interferon stimulated population (CD4_IFN_stim; *ISG15*, *IFIT3*, *TNF*, *IFI44*) consistent with a previous report^39^. Four DN populations were identified as double negative T cells (DN_T cell; *KANK1*, *NMB*, *KIAA0825*), double negative cytotoxic (DN_CTL; *ZEB2*, *NKG7*, *GZMB*, *KLRD1*, *CCL5*), gamma delta (δγ) T cells (gd_Tcell; *RHEX*, *GATA3*, LOC61156*5* (antigen *WC1.1-like*)) and MAIT-like cells (MAIT; *IL23R*, *RORC*, *KLRB1*). Comparison of the relative proportions of each population indicated only minor changes in cell frequencies when comparing NLC to dogs at each stage of disease progression.

**Figure 4.**
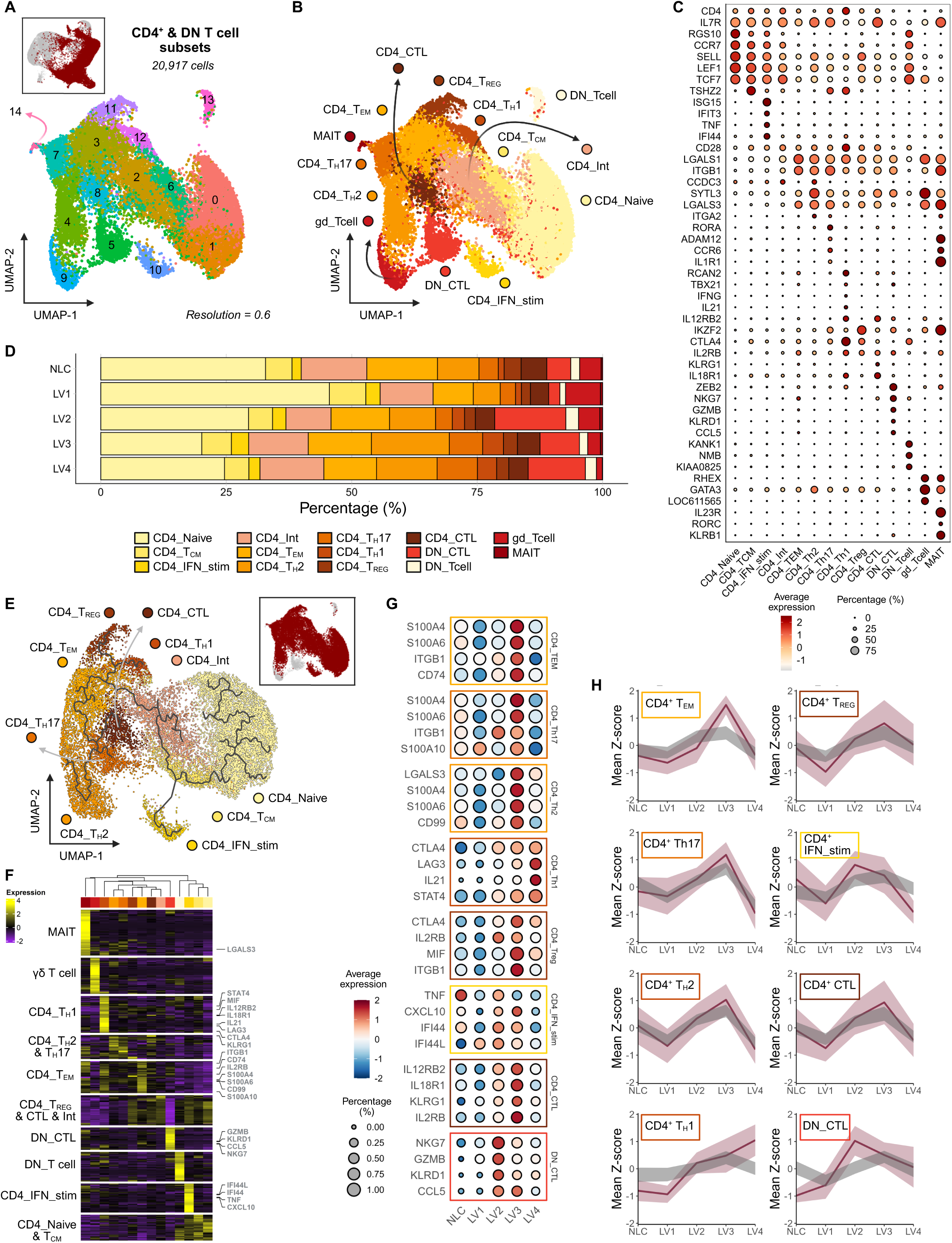
Peripheral CD4^+^ and DN T cells reveal a shift from naïve toward effector phenotypes. **(A)** UMAP visualization of 15 CD4^+^ and DN (double negative) T cell subclusters. The inset shows the projection of these cells onto the T/NK cell UMAP (Level 2). **(B)** UMAP visualization of 11 CD4^+^ and 4 DN annotated T cell populations. **(C)** Dot plot showing canonical marker gene expression and proportions in CD4^+^ & DN T cell subclusters (LOC611565: antigen *WC1.1*-like). **(D)** Bar plot depicting the proportion of cells across CanL clinical stages, colored by CD4^+^/DN cell population. See also Table S2. **(E)** Trajectory analysis of CD4^+^ T cell populations performed on a UMAP reduction of a CD4^+^ T cell population subset. **(F)** Heatmap visualization of top 50 differentially expressed genes (DEGs) of each subcluster. Showing DEGs and subclusters arranged by hierarchical clustering. Highlighted DEGs correspond to the those visualized in Figure 4G. See also Table S4. **(G)** Dot plots showing expression and proportions within specific cell populations across CanL clinical stage, showing representative DEGs of the subcluster’s DEG signature. **(H)** Line graphs of individual subclusters summarizing signature Z-scores across CanL clinical stages. The visualizations show the mean (dark burgundy line) and standard deviation (light burgundy ribbon) of Z-scores from the subclusters DEG signature (Top 50 DEGs), alongside a control (grey ribbon) of upper and lower confidence intervals from bootstrapping (n = 100,000) non-signature genes. See also Figure S4 and Table S4.

We confirmed differentiation of naïve CD4^+^ T cells into T_CM_ and through to T_EM_ T_H_1 and T_H_2 and T_REG_ cells using trajectory analysis (**Figure 4E)** and by hierarchical clustering of cluster associated differentially expressed genes (DEGs) (**Figure 4F and Table S4**). These analyses also indicated that CD4_IFN_stim likely originated from T_CM_ cells, as supported by expression of T_CM_ canonical markers (**Figure 4C**).

We next examined relative mRNA abundance for all signature genes that defined these T cell and DN cell populations (**Figure 4G**). Like B cells, we noted marked variability in the transcript abundance for population-defining signature genes across disease stage, leading us to develop a novel approach to visualize this data. We generated a Z-score for signature gene expression at each disease stage and accounted for stochastic variability in gene expression by bootstrapping with random gene pools (see **STAR**⍰**METHODS**). This analysis demonstrated that, regardless of changes in frequency of each subcluster between LV stages (**Figure 4D**), the transcript abundance for signature genes varied over disease course. Of note, transcripts for CD4_T_H_1 signature genes, included those associated with exhaustion (*LAG3*, *CTLA4*) were more highly abundant at LV4, the terminal stage of disease. In contrast, transcripts for signature genes associated with CD4_T_EM_, CD4_IFN_stim and DN_CTL were more abundant at either LV2 or LV3 (**Figures 4H**, **S4, and Table S5**), stages previously reported to be associated with the highest production of IFNγ and control of parasitemia^48,49^.

### CD8□ T cells

Unsupervised clustering of CD8^+^ and cytotoxic DN T/NK cells revealed 16 transcriptionally distinct subclusters (**Figure 5A-C and Table S4**). The relative abundance of each subset by disease class is shown (**Figure 5D** and **Table S2;** see **STAR**⍰**METHODS).**

**Figure 5.**
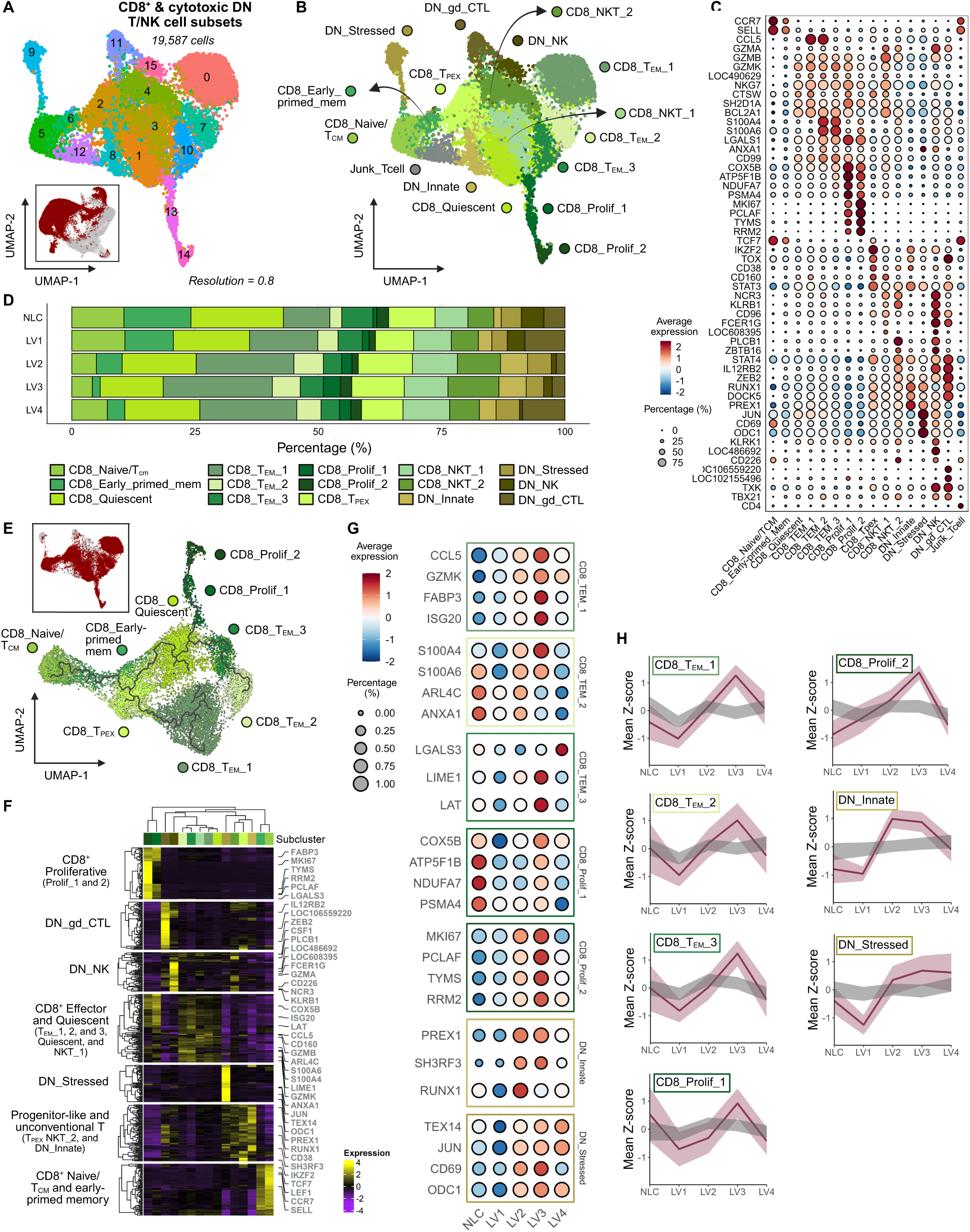
Divergent CD8□ T cell trajectories from a T_PEX_ state culminate in effector and proliferative programs at severe CanL. **(A)** UMAP visualization of 16 CD8^+^ and cytotoxic DN T/NK cell subclusters. The inset shows the projection of these cells onto the T/NK cell UMAP (Level 2). **(B)** UMAP visualization of 11 CD8^+^ and 4 cytotoxic DN annotated T cell populations. **(C)** Dot plot showing canonical marker expression and proportions of annotated CD8+ and cytotoxic DN T/NK cell subclusters (LOC486692: *NKG2A/NKG2B*-like; LOC490629: *GZMH*; LOC608395: NK-lysin-like; LOC106559220: *CMRF35*-like; LOC102155496: *TRDC*-like). **(D)** Bar plot depicting the proportion of cells across CanL clinical stages, colored by CD8^+^ & cytotoxic DN populations. See also Table S2. **(E)** Trajectory analysis of CD8^+^ T cell populations performed on a UMAP reduction of a CD8^+^ T cell population subset. **(F)** Heatmap visualization of top 50 differentially expressed genes (DEGs) of each subcluster. Showing DEGs and subclusters arranged by hierarchical clustering. Highlighted DEGs correspond to the those visualized in Figure 5G. See also Table S4. **(G)** Dot plots showing expression and proportions within specific cell populations across CanL clinical stage, showing representative DEGs of the subcluster’s DEG signature. **(H)** Line graphs of individual subclusters summarizing signature Z-scores across CanL clinical stages. The visualizations show the mean (dark burgundy line) and standard deviation (light burgundy ribbon) of Z-scores from the subclusters DEG signature (Top 50 DEGs), alongside a control (grey ribbon) of upper and lower confidence intervals from bootstrapping (*n* = 100,000) non-signature genes. See also Figure S5C and Table S4.

We identified two main CD8^+^ T cell differentiation trajectories (**Figure 5E-F**) originating from CD8_Naive/T_CM_ and CD8_Early-primed_mem cells (*CCR7*; *SELL*). Early-primed memory CD8□ T cells closely resembled naïve/T_CM_ cells in transcriptional profiles, sharing lymphoid-homing and survival markers (**Figure 5C** and **S4B**) but exhibiting higher expression of cytotoxic and effector genes (e.g., *GZMB*, *NKG7*, *ZEB2*) (**Figure S5A**). In the absence of detectable transcripts for PD-1, CD8_T_PEX_ cells (*n* = 1,895) were tentatively defined by *TCF7* along with high transcript abundance for genes associated with T cell dysfunction and progenitor exhaustion (*IKZF2*, *TOX*, *CD38*, *CD160*, and *LRBA*), immunoregulatory signaling (*STAT3*), and inhibition of TCR and cytokine signaling (*INPP5D* and *PLCL2*) (**Figure 5C and S4B**). One trajectory then progressed toward a cytotoxic effector phenotype (T_EM__1; *CCL5, GZMB, NKG7, SH2D1A, CTSW*). The other transitioned through a putative quiescent population (*BLC2A1*) before bifurcating into either i) proliferative cells encompassing CD8_Prolif_1 (with increased metabolism-associated transcripts including *ATP5F1B, COX5A, NDUFA7, FABP3,* as well as proteasome-related *PSMA4*) and CD8_Prolif_2, distinguished by higher expression of proliferation markers (*MKI67*, *PCLAF*, *TYMS*) and genes linked to DNA replication (*PCNA*, *RRM2*), DNA repair (*RAD51*), centrosome dynamics (*TPX2* and *SPC24*), and cell cycle progression (*BIRC5*, and *MYBL2*) but reduced metabolic activity, or ii) T_EM_ cells that transitioned from metabolically active (CD8_T_EM__3; *ATP5F1B* and *COX5B*) to a more migratory effector state (CD8_T_EM__2; *S100A4*, *S100A6*, *LGALS1*, *CD99*, and *ANXA1*). As with CD4^+^ T cells, we noted marked fluctuations in the average expression of signature genes (**Figure 5G**) associated with each CD8^+^ T cell subpopulation, with a general trend towards higher transcript abundance at LV3 for effector (T_EM__1, T_EM__2, T_EM__3) and proliferative (Prolif_2) populations (**Figure 5H and Table S5**). One cluster expressing *CD4* was determined to be misclassified (*Junk_Tcell*) (**Figure 5A-C**) and was removed from further analysis.

### CD8^+^ NKT cells and other innate lymphocytes

Of the two transcriptionally distinct CD8^+^ NKT-like populations identified, CD8_NKT_1 (*n* = 1,674 cells) was characterized by abundant transcripts for cytotoxic granule-associated (*GZMA*, *GZMB*, *GZMK*, *GZMH* (LOC490629), *NKG7, NK-lysin like* (LOC608395)), NK receptors (*CD160*, *KLRB1*, *NCR3, CD96*), and pro-inflammatory cytokines genes (*CSF1*, *IL2RB*) (**Figure 5C**). CD8_NKT_2 cells (*n* = 1,517 cells) had reduced transcripts for cytotoxic genes but were enriched for transcriptional regulators and cytokine signaling components (*ZBTB16*, *STAT4*, *IL12RB2*, *ZEB2, PLCB1*) suggesting a poised NKT-like population with lower effector commitment (**Figure 5C**).

DN_Innate T cells (*n* = 861 cells) exhibited a regulatory-like phenotype, marked by the expression of transcriptional and chromatin regulators (*IKZF2*, *RUNX1*, and *ZEB2*) alongside signaling and trafficking genes (*PREX1*), and low cytotoxic profile (**Figure 5C**). DN_Stressed T cells (*n* = 814 cells) had elevated transcripts for immediate early response genes (*FOSB* and *JUN*), cytokine mediators (i.e., *TNF* and *CD69*), and metabolic stress-related genes (*ODC1* and *MAT2A*). A minor DN_NK cell population (*n* = 316 cells) lacked CD3, CD4, or CD8 transcripts, but was enriched for cytotoxic NK-associated transcripts and innate immune effectors (including *NCR3* (NKp30), *KLRK1* (NKG2D), *NKG2A/NKG2B-like* (LOC486692), *KLRB1* (CD161), *CD226*, *IL12RB2*, *FCER1G*, *GZMA*, and *GZMB* (**Figure 5C** and **S4B**). DN_gd_CTL cells (*n* = 790 cells; CMRF35-like (LOC106559220) and TRDC-like (LOC102155496)) also had a cytotoxic transcriptional profile (*GZMA*, *TBX21,* and *TXK*) alongside signaling components such as *DOCK5* (**Figure 5C**). Among DN subsets, signature genes associated with DN_Innate and DN_Stressed populations were transcriptionally suppressed at LV1 but elevated at LV2 and LV3. Other populations displayed minimal changes in signature gene transcript abundance (**Figures 5H, S5C, and Table S5**).

### Myeloid cells

Myeloid cells are key host cells for *Leishmania* and play vital roles as immune effector cells and in immunoregulation^21,22^. Our analysis identified 16 populations of myeloid cells (**Figure 6A**), which were classified by canonical markers (**Figure 6B and C**). Monocytes (*n* = 11,586 cells; *LYZ, CSF1R, CTSS*) were the dominant cell type, with six sub-populations. Two could be annotated as classical (Mo_Classical: *FCGR1A* (CD64), *SELL* (CD62)); and non-classical monocytes (Mo_Non-Classical: LOC478984 (CD16a ortholog), *PECAM* (CD31), *n* = 1,978 cells), based on human and mouse nomenclature. The other four monocyte subsets (Mo_1, *n* = 2,851 cells; Mo_2, *n* = 2,821 cells; Mo_3, *n* = 1,891 cells; Mo_4, *n* = 501 cells) did not directly align with conventions commonly adopted for humans and mice. Neutrophils (*n* = 2,131 cells; *CSF3R, MEGF9, PADI4*), eosinophils (*n* = 685 cells; *ITGAM, CCR3, IL5RA*), and basophils (*n* = 364 cells; *CPA3, MS4A2, IL3RA*) were also identified. Three minor populations of dendritic cells (DCs) (*n* = 672 cells; *CD86, FLT3*) were found and further annotated as conventional DC type II (cDC2, *n* = 420 cells;*, PID1, CD2, CLEC12A*^50^), precursor DCs (preDCs, *n* = 144 cells; *IL3RA, PGLYRP2, IRF4*) and plasmacytoid DCs (pDCs, *n* = 98 cells; *IGF1, RARRES2, TLR7*^51^). Presumptive doublet populations of T cells (*CD3E,* LOC607937 *(TRAV9-2),* LOC480788 *(TRBC1)),* B cells (*CD79A, CD19, MS4A1*) and platelets (*PPBP, TUBB1, GP9*) were subsequently removed from further analysis.

**Figure 6.**
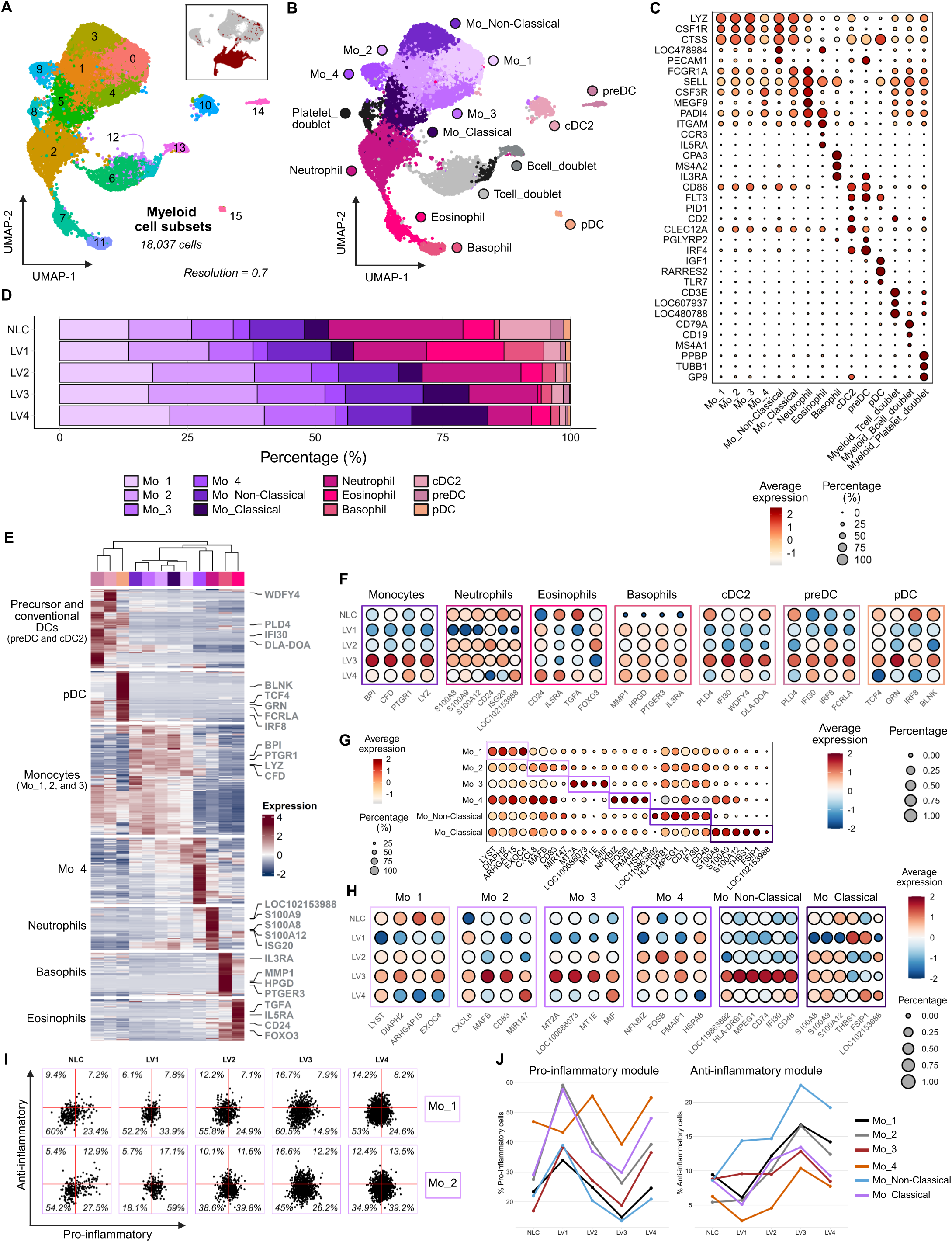
Myeloid compartment characterized by monocyte expansion and dynamic inflammatory monocyte profiles. **(A)** UMAP visualization of 16 myeloid subclusters. The inset shows the projection of these cells onto the PBMC UMAP (Level 1). **(B)** UMAP visualization of annotated myeloid cell populations. **(C)** Dot plot showing canonical marker gene expression and proportions in myeloid subclusters (LOC47894: *FCGR3A* (CD16a); LOC480788: *TRBC1*; LOC607937: *TRAV9-2*). **(D)** Bar plot depicting the proportion of cells across CanL clinical stages, colored by myeloid cell population. See also Table S2. **(E)** Heatmap visualization of the top 50 differentially expressed genes (DEGs) of each subcluster. Showing DEGs and subclusters arranged by hierarchical clustering. Highlighted DEGs correspond to those visualized in Figure 6F. See also Table S4. **(F)** Dot plots showing expression and proportions within specific cell populations across CanL clinical stage, showing representative DEGs of the subcluster’s DEG signature. **(G)** Dot plots showing expression and proportions within the six monocyte populations of representative DEGs of a monocyte specific DE analysis. **(H)** Dot plots showing expression and proportions within individual monocyte populations across CanL clinical stage, showing monocyte-specific DEGs also highlighted in Figure 6G. **(I)** Scatter plots of inflammatory profiling that juxtapose pro- and anti- inflammatory module scores of Mo_1 and Mo_2 cells at different clinical stages of CanL. Plots classify cells as pro-inflammatory (top left), anti-inflammatory (bottom right) non-inflammatory (bottom left) or mixed inflammatory (top right); and show percentage of cells in each quartile. See also Figure S6A. **(J)** Line graphs summarizing inflammatory profiling of monocyte populations across CanL clinical stage. The graphs visualize percentages of cells classified as pro- or anti-inflammatory for monocyte populations at different clinical stages of CanL. See also Table S6.

Monocytes increased in frequency across CanL progression with other cell types contracting to varying extents (**Figure 6D and Table S2**). We generated signatures based on the top 50 DEGs (**Table S4**) and conducted hierarchical clustering to confirm the cell typing and cluster relationships (**Figure 6E**). Like lymphocytes, transcript abundance for representative signature genes associated with all myeloid cell lineages varied across disease stage (**Figure 6F**).

We next sought to further characterize monocyte subsets, given their increasing relative abundance during disease progression (**Figure 1G**). We conducted a differential expression analysis between each monocyte subset (**Figure 6G and Table S5**). Mo_1 had transcripts for hallmark genes associated with phagocytosis and actin remodeling (*DIAPH2*, *ARHGAP15*), as well as vesicle and lysosomal trafficking (*EXOC4*, *LYST*); Mo_2 had transcripts for hallmark genes associated with positive (*CXCL8*) and negative (*MAFB*, *MIR147* and *CD83*) regulation of inflammation ; Mo_3 had a more unique signature including metallothionein genes (*MT2A*, LOC100686073 (*MT1*) and *MT1E*) and *MIF*, suggestive of a response to stress or reactive oxygen species, and Mo_4 had a signature indicative of a stress response (*HSPA8*) and apoptosis (*PMAIP1*) along with associated transcription factors (*NFKBIZ, FOSB*). Transcripts abundance for signature genes across all monocyte subsets trended upwards as disease progressed from LV1 to LV3 but were reduced at LV4. This was most notable for genes attributed to the inflammatory regulation of Mo_2, the metallothionein-stress response of Mo_3, and the antigen presenting cell functions of non-classical monocytes. This suggests further regulation of gene expression in monocytes at t end-stage disease (**Figure 6H**).

### Myeloid cells modulate inflammatory status though disease

Pro and anti-inflammatory balance is critical for *Leishmania* survival in myeloid cells and is indicative of underlying lymphocyte driven responses. To investigate this aspect of myeloid biology in more detail, we adopted a module-based approach using manually curated pro- and anti-inflammatory genes of known relevance in leishmaniasis, and that would allow us to profile each population on a single cell basis (**Table S6**; see **STAR**⍰**METHODS**). Changes in the proportion of cells adopting a pro- or anti-inflammatory profile were then visualized over disease stage (**Figures 6I, J,** and **S6A**).

Strikingly, the proportion of pro-inflammatory cells within each monocyte subset peaked early in disease, declined by LV2 or LV3 and rose again at LV4. In contrast, the anti-inflammatory cell frequency increased progressively to LV3, before declining at LV4. Other myeloid populations did not follow this defined pattern (**Figure S6B**), suggesting that monocytes were highly responsive to and helped shape a dynamic immune landscape associated with different stages of disease progression.

## DISCUSSION

Here, we have developed the LeishDog Atlas, a comprehensive single-cell transcriptomic atlas of canine peripheral blood immune cells in control and *L. infantum*-infected dogs. Profiling 68,323 cells across 16 PBMC samples, we annotated 47 distinct immune populations. An open-access GitHub repository accompanies the Atlas, providing a rich resource for researchers working on companion animal infectious diseases and for those using the dog as a model organism. Leveraging the LeishDog Atlas to gain insights into CanL, we highlight the cellular heterogeneity and dynamic immune programs that shape CanL progression, offering new perspectives on how immune responses may shift across clinical stages. Given the shared immunopathological features of CanL and human VL, these findings may also inform understanding of systemic immune dysregulation in human disease.

The canine immune system, and that of other companion animals, remains relatively poorly characterized compared to more established model organisms, typically rodents. As in other fields, the advent of scRNA-seq has provided new insights into the normal immune cell heterogeneity and function. However, canine single-cell datasets remain limited, particularly in the context of infectious diseases^39,40,52–54^. Ammons et al. generated an atlas of circulating leukocytes from healthy and osteosarcoma-affected dogs, providing important insight into baseline and tumor-associated immune states^39^. More recently, Manchester et al. revealed T cell heterogeneity within the duodenal mucosa of dogs with chronic inflammatory enteropathy relative to healthy controls^53^. While both studies addressed naturally occurring conditions, infection remains largely unexplored at single-cell resolution. The LeishDog Atlas extends current canine single-cell efforts by defining the immune landscape of *L. infantum*-infected dogs and serving as a reference for infection-driven immune responses. We also identified additional peripheral immune cell states, including MAIT-like cells, cytotoxic γδ T cells, T_PEX_, and two CD8^+^ NKT subsets in both control and infected animals. The potential role these populations in CanL, as in human VL, requires further investigation, particularly within parasitized tissues.

Single-cell reference datasets are also critical for spatial transcriptomic analyses, enabling cell-type deconvolution within complex tissue microenvironments^55^. However, most available reference datasets reflect normal physiology or a narrow range of disease states, potentially omitting cell phenotypes associated with distinct inflammatory conditions. Here, we chose to generate a scRNA-seq atlas integrating data from control dogs and from dogs representing well-characterized clinical stages of CanL, an exemplar of mononuclear cell-dominated, chronic systemic inflammatory disease. The immune cell classifications established in the LeishDog Atlas should therefore facilitate the interpretation of other canine infectious diseases, including tick-borne infections, leptospirosis, and histoplasmosis.

Whilst the study was not designed to provide definitive evidence for immune changes across CanL disease progression, it nevertheless provides several new insights into the dynamics and complexity of the response of dogs to *L. infantum* infection. First, we observed significant shifts in T cell and myeloid cell ratios during disease progression, being initially skewed towards a T cell dominance early in disease (LV1 and LV2) and then switching to become dominated by myeloid cells at LV3 and LV4. Subsequent analysis demonstrated that the latter reflected an increase in frequency of classical as well as non-classical monocytes, as well as 4 additional monocyte populations defined by distinct function-related transcriptional signatures. By generating a module score to reflect transcript abundance for cardinal pro- and anti-inflammatory genes, these monocytes were characterized at single cell resolution and the respective balance of pro and anti-inflammatory monocytes mapped over disease course. Whereas pro-inflammatory cells dominated early during infection, monocytes with anti-inflammatory potential emerged at LV3. This inflammatory monocyte profile aligns with expected changes in CanL, which are regularly depicted as an early T_H_1 response that shifts into a T_H_2 response and ends in a chronic phase^22^. Unexpectedly, we also observed a reversal of this trend at LV4, suggesting that terminal disease may be associated with a pro-inflammatory burst that contributes to clinical outcome but is unable to impact parasite burden. Although cause and effect cannot be directly attributed, it is of note that this terminal stage alteration in monocyte activation bias was paralleled by a rise in CD4^+^ T_H_1 activity and a decline in CD8^+^ T cell effector function (as inferred from transcript abundance).

Secondly, we observed striking changes in signature gene transcript abundance during disease progression. This was observed in many lymphocyte and myeloid cell populations and was largely independent of shifts in relative cell frequency. Two explanations can be put forward to this observation. For cells with cognate antigen receptors, increased transcription may be associated with antigen-driven activation. Addressing this possibility is challenging in the absence of tools to identify antigen-specific B and T cells by scRNA-seq. Alternatively, and more likely given the range of cell populations involved, increased transcription may reflect a more global effect driven by changes in the systemic inflammatory environment. Mechanistically, this may relate for example to alterations in systemic cytokine production or to metabolic changes, both known feature of CanL^20,22^. Additional studies will be required to determine whether this may also reflect epigenetic reprogramming of immune cells over progressive disease. For example, epigenetic modifications (including histone modification and DNA methylation) have been shown to be central to the establishment of T cell developmental trajectories^56,57^ and in systemic lupus erythematosus and rheumatoid arthritis, altered DNA methylation can lead to overexpression of genes and T cell hyperactivation^58,59^.

Finally, it is known that CD8^+^ T cells can play a dual role in CanL, contributing both to protective immunity and to disease progression^60,61^. Flow cytometry studies have identified increased frequencies of CD8^+^ lymphocytes as a hallmark of low parasite burden and subclinical disease. For example, CD8^+^ T cell frequencies were strongly correlated with low bone marrow parasitism in naturally-infected Brazilian dogs^24^. In addition, significant increases in CD8^+^ T cell frequencies and decreased CD4:CD8 ratios have been reported in *L. infantum*-infected dogs^62,63^. However, the transcriptional signatures associated with CD8^+^ T cells in CanL have not previously been reported.

During chronic infections, due to persistent antigen exposure and consequent inflammation, CD8^+^ T cells undergo successive inhibitory changes (“exhaustion”), which leads to progressive but potentially reversible loss of cytotoxic function and pathogen-specific responses^64^. Consistent with this model, Singh et al. also reported upregulation of exhaustion markers (LAG-3 and TIM-3) on CD8^+^ T cells in VL patients, along with elevated transcripts for cytolytic effector genes, including *GZMA*, *GZMB*, *GZMH* and *PRF1*^65^. Likewise, we have previously shown that prior to onset of clinical disease, circulating CD8^+^ T cells from asymptomatic *L. infantum*-infected dogs exhibit an exhaustion-like phenotype, characterized by elevated IL-10 production and PD-1 expression^30^. In the current study, we did not observe consistent upregulation of exhaustion-associated transcripts, even in dogs at the most advanced CanL stage. Based on available markers, we did, however, identify a putative CD8^+^ T_PEX_ population. In contrast to healthy human blood, where such cells are very rare (usually <1%^66^) we found CD8^+^ T_PEX_ at much higher frequency (∼10-15%) irrespective of infection status or disease stage. This may reflect a different threshold for T_PEX_ differentiation in dogs compared to humans or a greater degree of underlying “chronic” inflammation / infection. Trajectory analysis placed these cells on the pathway leading to the terminal differentiation of all other effector CD8^+^ populations, in keeping with progenitor potential.

The study has some limitations. First, being cross sectional in design, immune cell trajectories within individual dogs over time could not be studied. Together with the modest sample size and potential for immune modulation through tick-borne co-infections, conclusions regarding changes in cell distributions need to be treated cautiously. Second, our analysis examined B and T cells heterogeneity and function independently of antigen-specificity. This is not uncommon in studies of this type, especially where the antigen repertoire of the infectious agent is complex and diverse. Third, annotation of the dog genome remains limited and not all relevant transcripts (notably cytokines) were detectable. Hence for some immune subsets (e.g, CD8^+^ T_PEX_), it was not possible to definitively map cells based on their counterparts in humans and rodents or functional potential. Fourth, many key immune processes occur beyond gene expression levels, including chromatin remodeling, post-transcriptional regulation, and modifications. As a result, some cell states and/or functions may not be captured in our dataset, especially if they are primarily regulated through non-transcriptional mechanisms^67–69^. Finally, we fully acknowledge that follow-up protein-level validation and functional assays will be required to confirm some of the observations made. Nevertheless, our findings offer a valuable framework for developing and testing new hypotheses, which may guide future mechanistic studies, deepen our understanding of CanL and VL immunopathology, and facilitate the use of scRNA-seq as a tool to evaluate future immune-targeted therapies for CanL.

## RESOURCE AVAILABILITY

### Lead contact

Further information and requests for resources and reagents should be directed to and will be fulfilled by the lead contact, Paul M. Kaye.

### Materials availability

Materials and reagents generated in this study are available upon a reasonable request from the Lead Contact and may require a completed Material Transfer Agreement.

### Data and code availability

Transcriptomics data have been deposited at NCBI GEO (GSE313451) and will be publicly available as of the date of publication. All analysis scripts used for transcriptomic analyses are maintained in a private, free-of-cost access for reviewers GitHub repository (https://github.com/DanHolbrook/LeishDog_Atlas). Any additional information required to re-analyze the data reported in this paper is available from the Lead Contact upon request.

## ACKNOWLEDGEMENTS

This work was supported by the National Institutes of Health (NIH) under grant R01AI171971. PMK is also supported by a Wellcome Investigator Award (#224290). The authors thank the animal caretakers who participated in the study, Sally James and Lesley Gilbert (Biosciences Technology Facility, University of York) for technical assistance, and Nidhi Dey for advice on analytical approaches.

## AUTHOR CONTRIBUTIONS

Conceptualization: CAP, PMK; Data curation: DJH, DPU, KIC, MCW; Formal analysis: DJH, DPU; Funding acquisition: CAP, PMK; Investigation: DJH, DPU, MCW, NB; Methodology: DJH, DPU, JJO, SD; Project administration: CAP, PMK; Resources: CAP, PMK; Supervision: CAP, PMK; Validation: CAP, PMK; Visualization: DJH, DPU; Writing – original draft: DJH, DPU; Writing – review & editing: all authors.

## DECLARATION OF GENERATIVE AI AND AI-ASSISTED TECHNOLOGIES IN THE WRITING PROCESS

The authors used ChatGPT (OpenAI) for language editing and limited coding support during manuscript preparation. All content generated with this tool was reviewed and edited by the authors, who take full responsibility for the accuracy and integrity of the published work.

## SUPPLEMENTAL INFORMATION

Document S1. Figures S1-S6. Tables S1-S3, and supplemental reference.

Table S4. Differential expression analysis results. Related to Figures 2, 4, 5, 6, S3, S4, and S5.

Table S5. Average signature expression across CanL for CD4^+^ and CD8^+^ populations. Related to Figures 4 and 5.

Table S6. Curated genes used for inflammatory profiling of myeloid cells; genes expressed in 10% of any subcluster were selected and organized into pro- or anti-inflammatory. Related to Figure 6.

## STAR⍰METHODS EXPERIMENTAL MODEL

### Animals

All animal work was reviewed and approved by the University of Iowa Institutional Animal Care and Use Committee (IACUC). Of the 18 dogs included in this prospective cohort study, 15 were US-owned dogs (11 males and 4 females). Leishmaniasis, in all species, has been shown to have a sex bias^70^. Complete physical examinations were performed by licensed veterinarians following standard practices, and peripheral whole blood samples were collected once for *L. infantum* detection via quantitative PCR (qPCR), complete blood count, serum chemistry, and tick-borne disease serology. Thirteen of these dogs were naturally infected with *L. infantum*, while 2 served as non-*Leishmania*-infected controls (NLC). CanL clinical staging was determined using the LeishVet scoring system^20^. A pooled sample from 3 UK beagles of unknown age and sex (ENVIGO) was also collected and considered as one additional NLC case. Each individual dog was treated as an independent experimental unit and subsequently grouped for analysis according to *Leishmania* infection status and clinical stage. Among the naturally infected dogs, 2 were staged as LeishVet score 1 (LV1; mild), 6 LV2 (moderate), 3 LV3 (severe), and 2 LV4 cases (terminal). The demographics and clinical parameters of all dogs are described in **Table S1**. Dogs with severe and terminal disease (LV3/LV4) exhibited significantly lower red blood cell counts, hematocrit, and hemoglobin, a characteristic typically associated with late-stage CanL progression^71^.

No *a priori* sample size calculation was performed. This study was observational in nature and based on the availability of dogs presenting for veterinary examination. Therefore, sample size was determined by case availability during the study period rather than by a predefined hypothesis-driven primary outcome.

No formal study protocol was submitted prior to study initiation, as no interventions, experimental treatments, or randomization were used. Dogs included in this study were privately owned and maintained in their home environments prior to and at the time of sampling. Housing, diet, and husbandry conditions were not controlled as part of the study, and no additional monitoring beyond standard veterinary practice was required. No animals were excluded from the analysis after enrollment. No study-related adverse events were observed or expected, and no humane endpoints were defined, as animals were not subjected to experimental manipulation.

Clinical and laboratory personnel were aware of dog identities but not of final group allocation at the time of assessment, as LeishVet staging was determined following combined clinical evaluation as well as blood and biochemistry results. Bioinformatic and statistical analyses were conducted with full knowledge of group allocation.

## METHOD DETAILS

### PBMC isolation

Within 24 hours of collection, whole-blood samples were diluted in Dulbecco’s phosphate-buffered saline (DPBS 1X), underlaid with Ficoll–Paque Plus, and centrifuged (500g, 30 min, without brake). Immune cells at the interface were collected and washed in DPBS 1X. They were then resuspended in chilled RPMI-1640 with L-glutamine media, supplemented with 40% heat-inactivated fetal bovine serum (FBS), 1% MEM non-essential amino acids solution (100X), and 100μg/mL penicillin-streptomycin. Cells were counted via hemocytometer, centrifuged (300g, 10 min) and resuspended in CryoStor® CS10 (StemCell Technologies) for cryopreservation. Cryopreservation used a slow rate-controlled cooling method, within an isopropanol freezing container to freeze to -80°C, before transfer to liquid nitrogen. Later they were shipped to UoY (University of York, UK) and upon arrival stored in liquid nitrogen until preparation for scRNA-seq.

### Sample preparation for scRNA-seq

Samples were defrosted and diluted in warm RPM-1640 supplemented with 10% FBS, 100μg/mL penicillin-streptomycin and 1% L-glutamine. Cells were washed twice by centrifugation (300g, 10 minutes) and resuspension in the supplemented media, after which dead cells were removed using a Dead Cell Removal Kit (Miltenyi Biotec) according to the manufacturer’s instructions. Remaining cells were adjusted to approx. 40,000 cells/mL in 0.04% ULTRA pure Bovine Serum Albumin. Library preparation was performed by the Genomics Laboratory, Biosciences Technology Facility, UoY using Chromium Next GEM Single Cell 3’ Kit v 3.1 (10X Genomics). Sequencing was performed by Illumina using the NovaSeq 6000 platform.

### Processing scRNA-seq data

FASTQ files were processed using Cell Ranger (v7.2.0, 10x Genomics) and aligned against the canine reference Dog10K_Boxer_Tasha (canFam6) from NCBI (GCF_000002285.5).

Quality control was performed using Cell Ranger’s QC algorithm, Seurat (v5.0.3)^72^ pipeline, and R (v4.2.1). The manual filtering of cells was conducted using the QC metrics: *nCount_RNA*, *nFeature_RNA*, *Log10GenesPerUMI* and *percent.mt*. For *nCount_RNA* and *nFeature_RNA*, a sample specific approach was used due to inter-sample variation. Cells with a *Log10GenesPerUMI* (log10(*nFeature_RNA*) / log10(*nCount_RNA*)) of >0.83, and mitochondrial/total RNA expression (*percent.mt*) of <5% were excluded. Manually selected of mitochondrial genes were used to derive percent.mt due to inadequate labelling of the reference gtf.

### PBMC Integration and clustering (*Level 1*)

A total of 68,323 cells were integrated into a PBMC Seurat object from 16 samples using a Seurat pipeline. Cells were normalizated (*SCTransform* function) using *glmGamPoi*-based regression of *nCount_RNA*, *nFeature_RNA*, *percent.mt* and *percent.rb* (percentage of counts from ribosomal genes). Cells were integrated (per sample) using anchors via 2000 integration features. Dimensionality reduction (*RunPCA* function) was performed using principal component analysis (PCA), and the top 30 principal components were selected via graph-based clustering. This was imputed using nearest neighbour identification (*FindNeighbors* function) and K-means clustering (*FindClusters* function). Clustering of 0.3 resolution was selected based on analysis using *clustree* (v0.4.4)^73^, and UMAP (Uniform Manifold Approximation and Projection) (*RunUMAP* function) was run on the top 30 PCA (npcs) as input to visualize the cells in a 2-dimensional space.

Analysis of major cell types in the PBMC dataset (resolution 0.3) indicated that cluster #9, although initially classified as T cell, also contained plasma cells. Subsequent higher-resolution clustering (0.8) segregated these populations, revealing cluster #25 as distinct plasma cell population, which was then manually re-assigned as B cell.

### Subcluster re-integration pipeline (*Level 2 and 3*)

Subclustering of major cell groups (T cells, B cells and Myeloid) was conducted by subsetting the appropriate clusters and splitting the data per sample (*SplitObject* function) to facilitate re-integration. Integration pipeline (as in in subsection *PBMC Integration and Clustering*) was used for sub-clustering but with specific npcs and resolution per cell type (Tcell – 20 dims, res: 2.0; Bcell – 20 dims, res: 0.7; Myeloid – 20 dims, res: 0.7). As T cells play central role in VL immunity, higher-resolution subclustering (2.0) was required to resolve the underlying heterogeneity within this compartment and to clearly delineate functionally distinct populations.

A third level of subclustering further split the T cells into CD4/DN (CD4^+^ T cells and double negative T cells) and CD8/Cyto_DN (CD8^+^ T cells and cytototic DN cells) using abovementioned methodology with specific input PCA dimensions and resolution (CD4/DN, npcs: 20, res: 0.6; CD8/Cyto_DN, npcs: 20, res: 0.8).Additionally, the CD4/DN cells were integrating with 4000 integration features to increase the anchor points of all cells to account for lower number of variable features within these cells.

Finally, the uncharacterized gene (LOC119871164), although variable in the dataset was removed as a variable feature for CD4/DN. As this single gene disproportionately impacted clustering, generating artifactual clusters.

### Cell type annotation

Expression of canonical markers were used to manually annotate the datasets. The approach used existing canine datasets and atlases, along with human and mice markers believed to be orthologous. Canonical marker expression was visualized by dot plots (**Figure 1D**, **2C**, **3C**, **4C**, **5C** and **6C**). Major cell types of the PBMC (*L*evel 1) object were further visualized by feature plots of module scores (*AddModuleScore* function) of the canonical markers (Figure 1C).

To validate our manual cell type annotation of PBMC clusters, we performed reference mapping using the Azimuth^34^. To project our dataset onto a well-curated, pre-annotated reference atlas to obtain predicted cell identities based on transcriptomic similarity. We applied the *RunAzimuth* function in Seurat (v5.0.3) using the human PBMC reference dataset from the Azimuth database (https://azimuth.hubmapconsortium.org/). The Azimuth algorithm identified anchors between the reference and query datasets, transferring reference-derived labels to each cell in our dataset based on shared transcriptional signatures. Predicted cell annotations were obtained at the *Level 1* resolution and visualized using a UMAP embedding generated from our dataset (**Figure 1F**).

### Doublet identification and removal

Doublet removal relied on identification through canonical markers. At *Level 2* subclustering, which separated the major cell groups (B cells, T cells and Myeloid), doublet populations were identified by co-expression of canonical markers of two cell groups. For instance, subcluster 6 (sc6) and sc12 of the Myeloid object showed co-expression of monocyte/neutrophil (*LYZ, CSF1R*, *CTSS*, *PADI4*, *CSF3R*, *MEGF9*) and T cell (*CD3E*, *LOC607937*, *LOC480788*) markers (**Figure S2A**). Likewise, Myeloid-Bcell_doublet (sc13) and Myeloid-platelet_doublet (sc8) populations were found with B cell (*CD79A*, *CD19*, *MS4A1*) or platelet (*PPBP*, *TEBB1*, *GP9*) markers. Using equivalent methodology in other objects, B_Tcell_doublet (sc7) and B_Myeloid_doublet (sc9) populations were found in the B cell object (**Figure S2B**), whereas only a single T_Myeloid_doublet (sc11) population was found in the T cell object (**Figure S2C**).

Further confirmation of canonical marker co-expression in doublet populations was confirmed by distinguishing doublets in the PBMC (**Figure S2D**) object via overlaying metadata from the *Level 2* subcluster objects into the *Level 1* object. This facilitated an overall comparison of marker expression in doublet and singlet populations (**Figure S2D**).

Populations identified as doublets were left in the datasets for completeness and visualization of the populations (**Figures 2A-C**, **3A-C**, and **6A-C**). After which they were removed prior to further analysis and use of the object (**Figures 2D-G**, **3D-E**, and **6D-J**). Similarly, the Junk_Tcell population of the CD8/Cyto_DN (*Level 3*) object was left in for visualization (**Figures 5A-C**) and removed prior to analysis and use of the object (**Figures 5D-H**).

## QUANTIFICATION AND STATISTICAL ANALYSIS

The primary outcome measure was the single-cell transcriptomic profile of peripheral blood mononuclear cells, assessed by scRNA-seq. Derived outcome measures included cell type and state annotation, relative cell population abundances, and differential gene expression associated with *Leishmania* infection status and clinical disease stage.

Statistical analyses were performed in R using established packages, including mixed-effects logistic regression, differential expression analysis, trajectory inference, module scoring, and pathway enrichment (see **Key Resources Table**). All analyses were conducted using the integrated dataset comprising samples from all dogs included in the study. Statistical methods appropriate for sparse, non-normally distributed single-cell transcriptomic data were used throughout, and model fit and diagnostics were evaluated using standard procedures implemented in the respective R packages where applicable.

### Mixed-effects logistic regression analysis

To assess alterations in immune cell composition across clinical stages, we applied a generalized linear mixed-effects model (GLMM), similar to that described by Fonseka et al.^42^ and Alladina et al^43^. We used the *glmer* function from the *lme4* R package (v1.1-35.5)^74^ to fit the following model:

*cbind(frequency, total_freq - frequency) ∼ 1 + LV.stage * cellgroup + (1 | orig.ident)*

Here, *frequency* represents the number of cells of a given type within each sample (*orig.ident*); *total_freq* is the total number of cells in that sample; *LV.stage* is a factor with five levels (NLC, LV1–LV4) representing clinical stage; and *cellgroup* (or *Tcell_celltype* in the T/NK cell subset analysis) indicates the major immune cell type. The term (1 | *orig.ident*) specifies a random intercept accounting for inter-sample variability. The model tested the interaction between clinical stage and cell type to determine whether the relative abundance of one cell population increased or decreased in relation to another across LeishVet-defined disease stages (LV1–LV4), using uninfected controls (NLC) as reference.

In our datasets, the estimated variance of the random intercept approached zero (singular fit), indicating minimal between-sample variation after accounting for fixed effects. Consequently, the model behaves equivalently to a fixed-effects binomial GLM, and coefficients were interpreted as population-level effects. Estimated marginal means were computed using the *emmeans* package (v1.10.6)^75^, and pairwise contrasts between cell groups were calculated within each clinical stage to derive log odds ratios (logOR) and 95% confidence intervals (**Table S3**). Fold-change in logOR relative to NLC was visualized as forest plots (**Figures 1 and 3**; see also **Table S3**). *P*-values were adjusted for multiple testing using the Benjamini–Hochberg false discovery rate (FDR) method.

### DEG signature Analysis

Differential expression (DE) analysis was performed using MAST (v1.27.1)^76^ within Seurat (*FindAllMarkers* function; *avg_log2FC* > 0 and *p_val_adj* < 0.05). Following this, the top 50 DEGs as ranked by significance (*p_val_adj*) were then used to define the gene signature per cell type.

Analysis of these signatures utilized heatmaps to provide further insight into differences between subclusters. *ComplexHeatmap* (v2.20.0)^77^ (*Heatmap* function) was used to visualize relative Z-score differences between subclusters and to perform hierarchical Euclidean clustering of the subcluster signatures to arrange both subclusters and genes. Resulting relationships are visualized in accompanying dendrograms. DEG signatures were further used to assess variation in disease progression within the individual subcluster as characterized by the signature. Relative change in expression across the disease were assessed on subsetted subclusters via dot plots that visualized Z-scores of the DEGs. For CD4/DN and CD8/NK objects this analysis was also summarized in line and ribbon plots, respectively visualizing the mean and standard deviation of the whole signature. These were calculated using DEG expression data (*AverageExpression* function) across CanL clinical stage within the subcluster subset. These results were scaled to Z-scores and used for means and standard deviations (*sd* function) of the 50 DEGs across each LV stage. This analysis was benchmarked against a bootstrap control of 50 non-signature genes with non-zero expression, and permuted 100,000 times with replacement. The bootstrapped control demonstrated as upper and lower confidence intervals of 95% (quantile function) was displayed as an additional grey ribbon.

### Trajectory Analysis

Trajectory analysis used to study differentiation of CD4^+^ and CD8^+^ T cells, was conducted using RStudio (v2024.09.1+394) with a newer version of Seurat (v5.2.1) and R (v4.4.2) (details of which are available on GitHub repository). Appropriate subclusters from the CD4 and CD8 dataset objects were subset and UMAP was re-run. This further analysis occurred in Monocle3 (v1.3.7)^78^, initially by converting the datasets from Seurat objects into cds objects (*as.cell_data_set* function) and then manually overlaying the subcluster metadata and UMAP embeddings into the cds objects. Trajectory analysis (*learn_graph* function) was calculated with normal pre-sets except for no closed loops.

### Myeloid Inflammatory Profiling Analysis

Inflammatory profiling of myeloid cells was conducted using a curated panel of inflammatory markers. The markers were filtered to retain only those expressed in ≥10% of cells within at least one myeloid subcluster. This unbiased filtering removed background signal of low frequency markers and improved specificity to the dataset and relevance to the canine system. Retained markers were subsequently classified as pro-inflammatory or anti-inflammatory and used for module score analysis (*AddModuleScore* function), scoring each cell simultaneously by both pro-inflammatory module and anti-inflammatory module. These scores were visualized as scatter plots at each clinical stage of CanL to demonstrate population changes in the frequency of cells considered pro-inflammatory, anti-inflammatory, non-inflammatory or mixed-inflammatory. Further line graph visualizations used the proportion of cells classified as either pro-inflammatory or anti-inflammatory to track changes to the inflammatory profile of myeloid populations over the progression of CanL.

### Gene ontology analysis

To further investigate the biological differences between two class-switched B cell clusters (B_1 and B_2), we performed DE analysis followed by pathway enrichment using Metascape^79^ (https://metascape.org). DE analysis was conducted in Seurat (v5.0.3) using the *FindMarkers()* function with subcluster B_2 against B_1. Genes were classified as up- or downregulated based on an adjusted *p* < 0.01 and |log□FC| > 0.5 threshold. The resulting gene sets were further refined by including only those with ≥ 10% expression difference (*pct.diff*) between clusters. The lists of significantly up-(B_2) and downregulated (B_1) genes were submitted separately to Metascape for *Gene Ontology (GO) Biological Process* over-representation analysis. Enrichment results (GO terms, enrichment score, *q*-value, and gene ratio) were exported and visualized in the R environment. The top 20 GO terms per cluster were ranked by adjusted *q*-value and enrichment score to highlight distinct functional programs between the two populations (**Figure S3B-C**).

## KEY RESOURCES TABLE

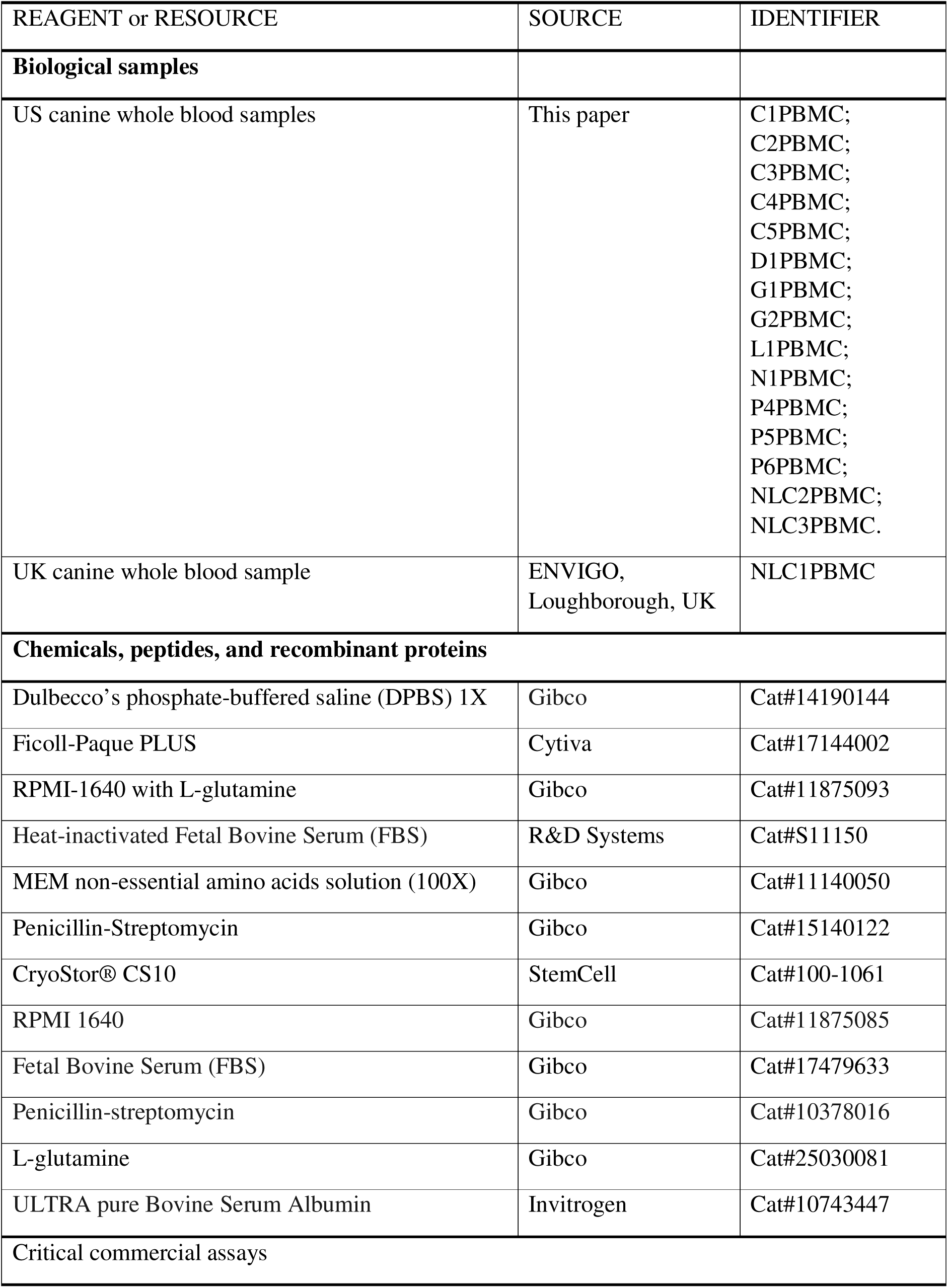
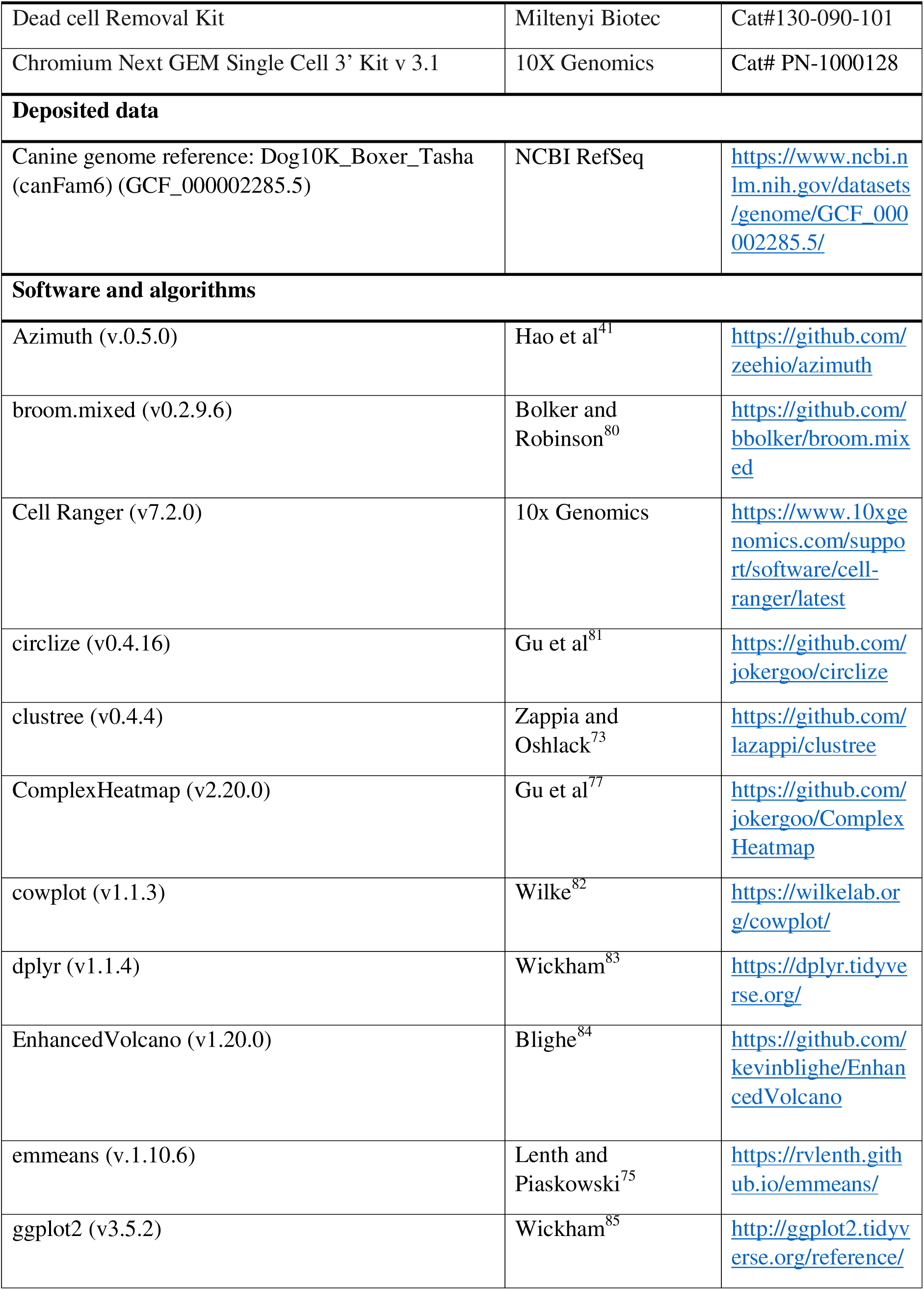
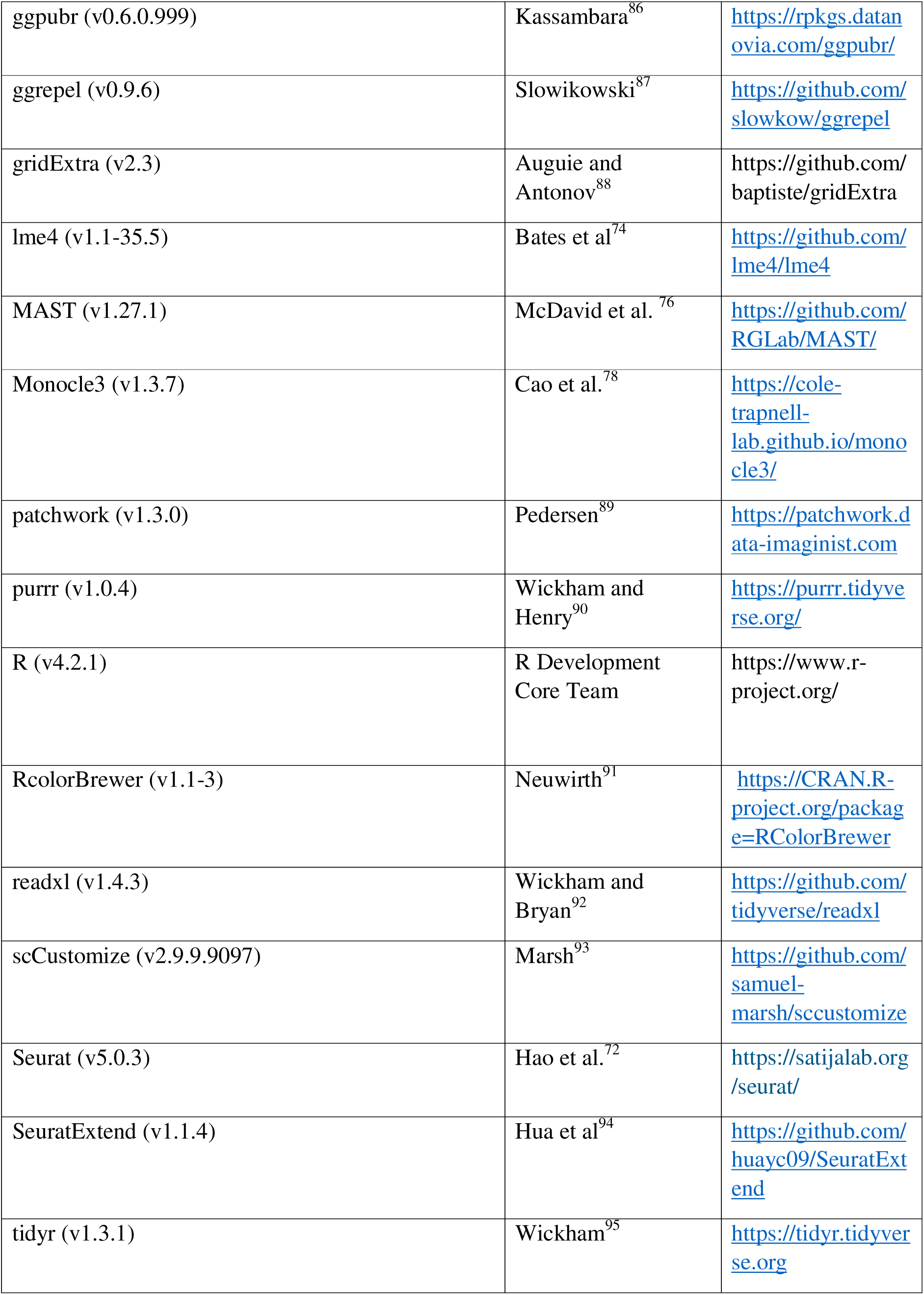
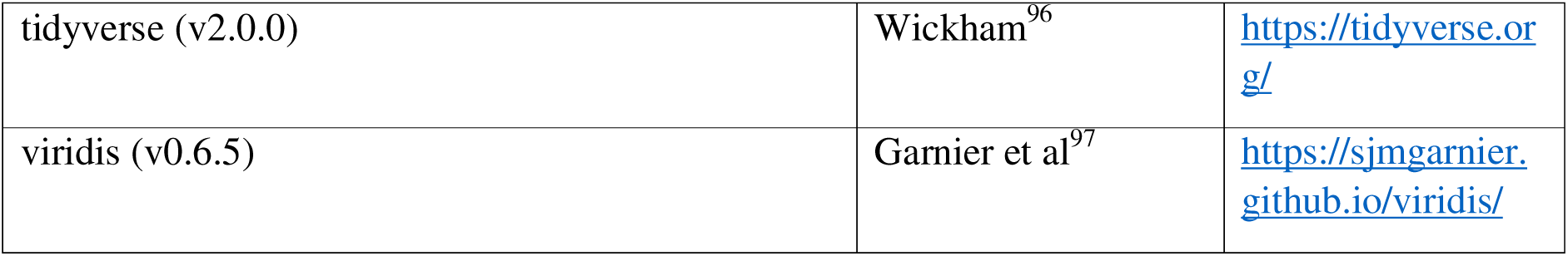

