## Supplementary Figures S1-S6 and Tables S1-S3 for "A single-cell transcriptomic atlas of peripheral blood immune cells spanning progressive canine leishmaniosis"

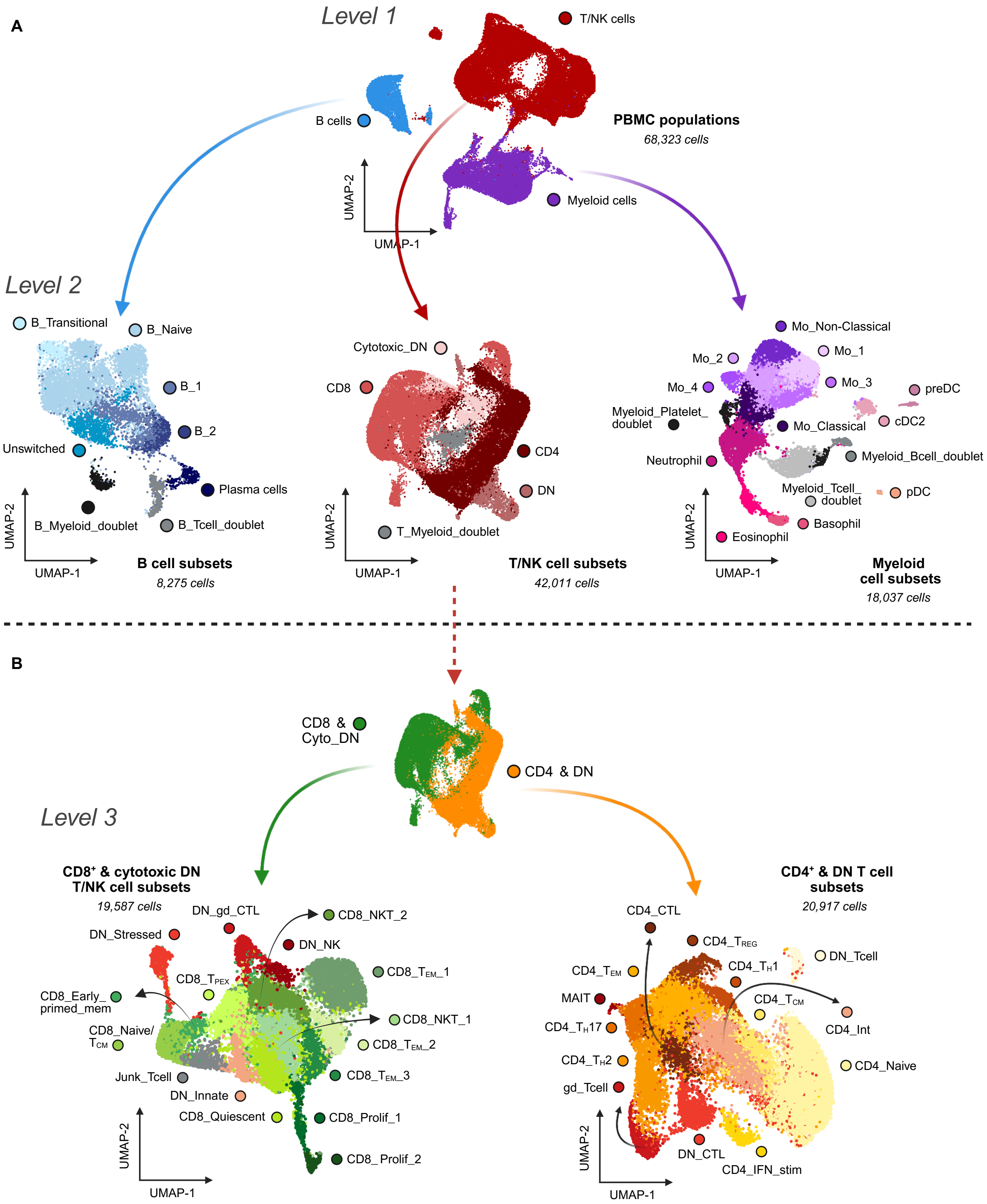

**Figure S1: Strategy for stepwise immune subsetting.**
Schematic of the subsetting strategy used to resolve major PBMC populations into Level 2 B, T/NK, and Myeloid subsets **(A)**; and T/NK cell populations into Level 3 CD4+ & DN and CD8+ & Cytotoxic DN subpopulations **(B)**.

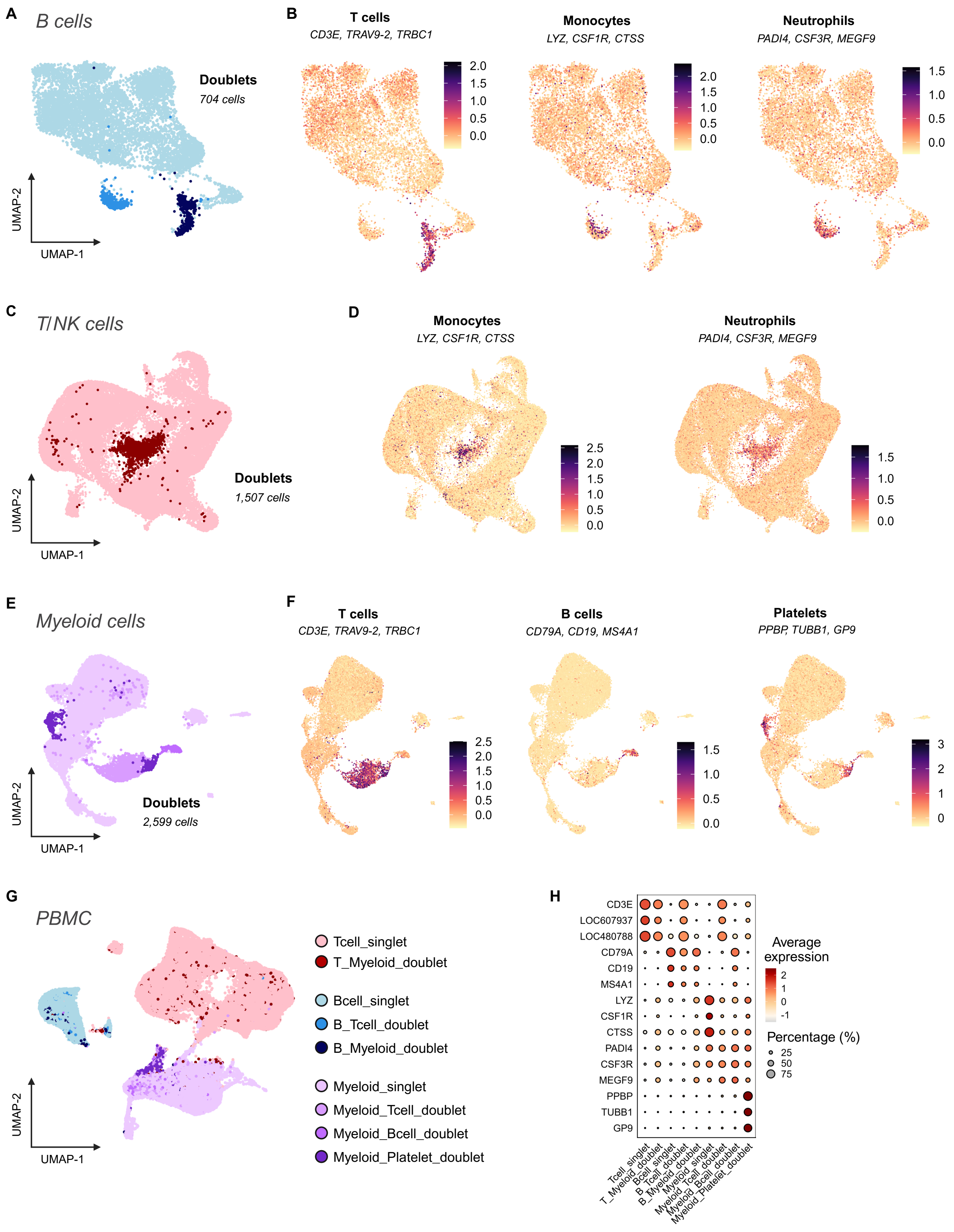

A
B cells
Doublets
704 cells
UMAP-2
UMAP-1
B
T cells
CD3E, TRAV9-2, TRBC1
Monocytes
LYZ, CSF1R, CTSS
Neutrophils
PADI4, CSF3R, MEGF9
C
T/NK cells
Doublets
1,507 cells
D
Monocytes
LYZ, CSF1R, CTSS
Neutrophils
PADI4, CSF3R, MEGF9
E
Myeloid cells
Doublets
2,599 cells
F
T cells
CD3E, TRAV9-2, TRBC1
B cells
CD79A, CD19, MS4A1
Platelets
PPBP, TUBB1, GP9
G
PBMC
Tcell_singlet
T_Myeloid_doublet
Bcell_singlet
B_Tcell_doublet
B_Myeloid_doublet
Myeloid_singlet
Myeloid_Tcell_doublet
Myeloid_Bcell_doublet
Myeloid_Platelet_doublet
H
CD3E
LOC607937
LOC480788
CD79A
CD19
MS4A1
LYZ
CSF1R
CTSS
PADI4
CSF3R
MEGF9
PPBP
TUBB1
GP9
Average expression
Percentage (%)
25
50
75

**Figure S2: Co-expression of dual canonical markers for doublet identification.**
**(A)** UMAP visualization showing B cell singlet and doublet populations within the Bcell object (see also Figure 2).
**(B)** Feature plots showing module scores for selected canonical marker genes used to identify and distinguish T cell (CD3E, LOC607937 (*TRAV9-1*); LOC480788 (*TRBC1*)), Monocyte (*LYZ*, *CSF1R*, *CTSS*) and Neutrophil (*PADI4*, *CSF3R*, *MEGF9*) populations in the Bcell object.
**(C)** UMAP visualization showing T cell singlet and doublet populations within the Tcell object (see also Figure 3).
**(D)** Feature plots showing module scores for selected canonical marker genes used to identify and distinguish Monocyte (*LYZ*, *CSF1R*, *CTSS*) and Neutrophil (*PADI4*, *CSF3R*, *MEGF9*) populations in the Tcell object.
**(E)** UMAP visualization showing Myeloid singlet and doublet populations within the myeloid object (see also Figure 6).
**(F)** Feature plots showing module scores for selected canonical marker genes used to identify and distinguish T cell (*CD3E*, *LOC607937* (*TRAV9*-1); *LOC480788* (*TRBC1*)), B cell (*CD79A*, *CD19*, *MS4A1*) and Platelet (*PPBP*, *TUBB1*, *GP9*) populations in the Myeloid object.
(**G**) UMAP visualization of the PBMC object displaying cells of the singlet and doublet populations. These were overlayed from the metadata of the level 2 (Myeloid, Tcell and Bcell) objects into level 1 (PBMC).
(**H**) Dot plot showing proportion and expression of canonical markers in singlet and doublet populations of the PBMC object generated as described in **B**, **D**, and **F**.

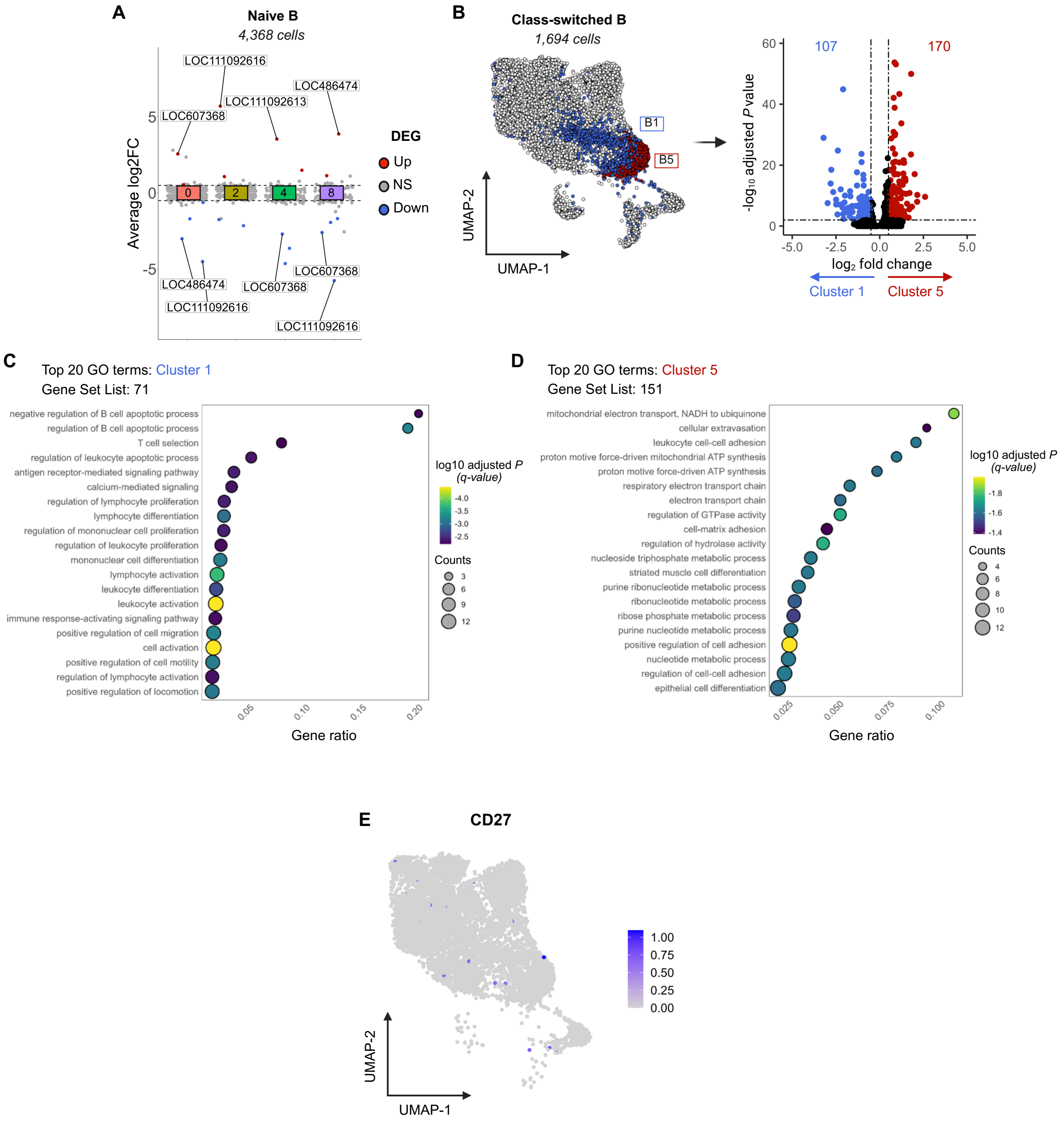

**Figure S3: Transcriptional and functional divergence among similarly annotated B cell subsets. Related to Figure 2.**
**(A)** Naive B cell subsets show minimal transcriptional differences and segregate according to the expression of specific immunoglobulin light chain genes. DEGs were defined by $|\log_2FC| > 0.5$ and adjusted $P < 0.01$. DEG: differentially expressed genes; LOC486474: *IGLC7*-like; LOC607368: *IGLV3-1*; LOC111092616 and LOC111092613: *IGLL1*-like; NS: non-significant.
**(B)** UMAP and volcano plot highlighting transcriptional divergence between class-switched B cell subsets (1 (B_1) and 5 (B_2)). DEGs were defined by $|\log_2FC| > 0.5$ and adjusted $P < 0.01$.
**(C and D)** Metascape[1] overrepresentation GO analysis of up- and downregulated genes between class-switched B cell subsets (5 vs 1). Downregulated genes (cluster 1, **(C)**) and upregulated genes (cluster 5 **(D)**) were analyzed for Biological Process enrichment ($|\log_2FC| > 0.5$, adjusted $p < 0.01$), with the top 20 terms displayed by gene ratio and log10 adjusted *P*-value.
**(E)** Feature plot showing minimal *CD27* expression across major B cell subpopulations.

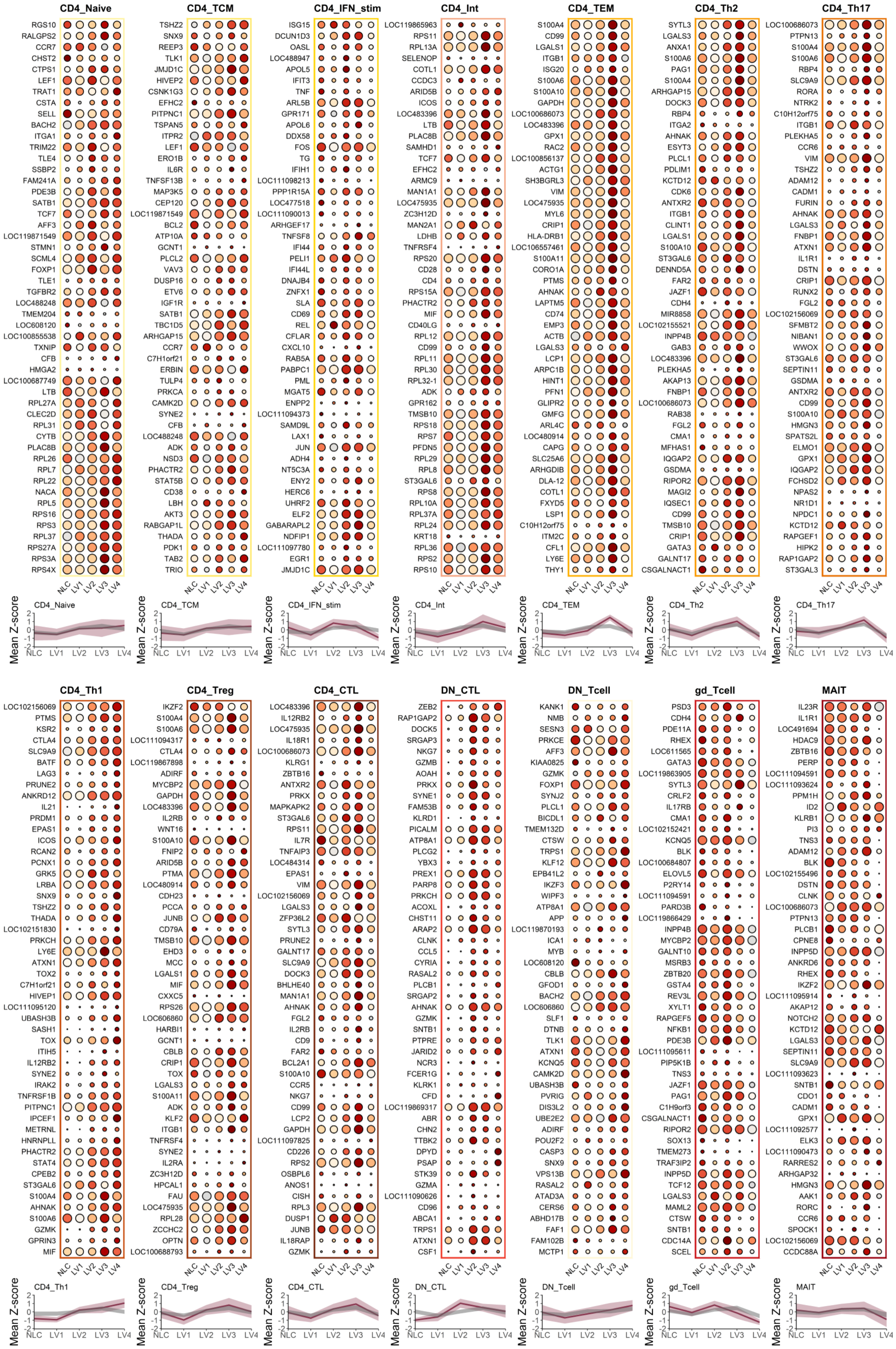

CD4_Naive
RGS10
RALGPS2
CCR7
CHST2
CTPS1
LEF1
TRAT1
CSTA
SELL
BACH2
ITGA1
TRIM22
TLE4
SSBP2
FAM241A
PDE3B
SATB1
TCF7
AFF3
LOC119871549
STMN1
SCML4
FOXP1
TLE1
TGFBR2
LOC488248
TMEM204
LOC608120
LOC100855538
TXNIP
CFB
HMGA2
LOC100687749
LTB
RPL27A
CLEC2D
RPL31
CYTB
PLAC8B
RPL26
RPL7
RPL22
NACA
RPL5
RPS16
RPS3
RPL37
RPS27A
RPS3A
RPS4X
CD4_TCM
TSHZ2
SNX9
REEP3
TLK1
JMJD1C
HIVEP2
CSNK1G3
EFHC2
PITPNC1
TSPAN5
ITPR2
LEF1
ERO1B
IL6R
TNFSF13B
MAP3K5
CEP120
LOC119871549
BCL2
ATP10A
GCNT1
PLCL2
VAV3
DUSP16
ETV6
IGF1R
SATB1
TBC1D5
ARHGAP15
CCR7
C7H1orf21
ERBIN
TULP4
PRKCA
CAMK2D
SYNE2
CFB
LOC488248
ADK
NSD3
PHACTR2
STAT5B
CD38
LBH
AKT3
RABGAP1L
THADA
PDK1
TAB2
TRIO
CD4_IFN_stim
ISG15
DCUN1D3
OASL
LOC488947
APOL5
IFIT3
TNF
ARL5B
GPR171
APOL6
DDX58
FOS
TG
IFIH1
LOC111098213
PPP1R15A
LOC477518
LOC111090013
ARHGEF17
TNFSF8
IFI44
PELI1
IFI44L
DNAJB4
ZNFX1
SLA
CD69
REL
CFLAR
CXCL10
RAB5A
PABPC1
PML
MGAT5
ENPP2
LOC111094373
SAMD9L
LAX1
JUN
ADH4
NT5C3A
ENY2
HERC6
UHRF2
ELF2
GABARAPL2
NDFIP1
LOC111097780
EGR1
JMJD1C
CD4_Int
LOC119865963
RPS11
RPL13A
SELENOP
COTL1
CCDC3
ARID5B
ICOS
LOC483396
LTB
PLAC8B
SAMHD1
TCF7
EFHC2
ARMC9
MAN1A1
LOC475935
ZC3H12D
MAN2A1
LDHB
TNFRSF4
RPS20
CD28
CD4
RPS15A
PHACTR2
MIF
CD40LG
RPL12
CD99
RPL11
RPL30
RPL32-1
ADK
GPR162
TMSB10
RPS18
RPS7
PFDN5
RPL29
RPL8
ST3GAL6
RPS8
RPL10A
RPL37A
RPL24
KRT18
RPL36
RPS2
RPS10
CD4_TEM
S100A4
CD99
LGALS1
ITGB1
ISG20
S100A6
S100A10
GAPDH
LOC100686073
LOC483396
GPX1
RAC2
LOC100856137
ACTG1
SH3BGRL3
VIM
LOC475935
MYL6
CRIP1
HLA-DRB1
LOC106557461
S100A11
CORO1A
PTMS
AHNAK
LAPTM5
CD74
EMP3
ACTB
LGALS3
LCP1
ARPC1B
HINT1
PFN1
GLIPR2
GMFG
ARL4C
LOC480914
CAPG
SLC25A6
ARHGDIB
DLA-12
COTL1
FXYD5
LSP1
C10H12orf75
ITM2C
CFL1
LY6E
THY1
CD4_Th2
SYTL3
LGALS3
ANXA1
S100A6
PAG1
S100A4
ARHGAP15
DOCK3
RBP4
ITGA2
AHNAK
ESYT3
PLCL1
PDLIM1
KCTD12
CDK6
ANTXR2
ITGB1
CLINT1
LGALS1
S100A10
ST3GAL6
DENND5A
FAR2
JAZF1
CDH4
MIR8858
LOC102155521
INPP4B
GAB3
LOC483396
PLEKHA5
AKAP13
FNBP1
LOC100686073
RAB38
FGL2
CMA1
MFHAS1
IQGAP2
GSDMA
RIPOR2
MAGI2
IQSEC1
CD99
TMSB10
CRIP1
GATA3
GALNT17
CSGALNACT1
CD4_Th17
LOC100686073
PTPN13
S100A4
S100A6
RBP4
SLC9A9
RORA
NTRK2
C10H12orf75
ITGB1
PLEKHA5
CCR6
VIM
TSHZ2
ADAM12
CADM1
FURIN
AHNAK
LGALS3
FNBP1
ATXN1
IL1R1
DSTN
CRIP1
RUNX2
FGL2
LOC102156069
SFMBT2
NIBAN1
WWOX
ST3GAL6
SEPTIN11
GSDMA
ANTXR2
CD99
S100A10
HMGN3
SPATS2L
ELMO1
GPX1
IQGAP2
FCHSD2
NPAS2
NR1D1
NPDC1
KCTD12
RAPGEF1
HIPK2
RAP1GAP2
ST3GAL3
NLC
LV1
LV2
LV3
LV4
Mean Z-score
2
1
0
-1
-2
CD4_Naive
CD4_TCM
CD4_IFN_stim
CD4_Int
CD4_TEM
CD4_Th2
CD4_Th17
CD4_Th1
LOC102156069
PTMS
KSR2
CTLA4
SLC9A9
BATF
LAG3
PRUNE2
ANKRD12
IL21
PRDM1
EPAS1
ICOS
RCAN2
PCNX1
GRK5
LRBA
SNX9
TSHZ2
THADA
LOC102151830
PRKCH
LY6E
ATXN1
TOX2
C7H1orf21
HIVEP1
LOC111095120
UBASH3B
SASH1
TOX
ITIH5
IL12RB2
SYNE2
IRAK2
TNFRSF1B
PITPNC1
IPCEF1
METRNL
HNRNPLL
PHACTR2
STAT4
CPEB2
ST3GAL6
S100A4
AHNAK
S100A6
GZMK
GPRIN3
MIF
CD4_Treg
IKZF2
S100A4
S100A6
LOC111094317
CTLA4
LOC119867898
ADIRF
MYCBP2
GAPDH
LOC483396
IL2RB
WNT16
S100A10
FNIP2
ARID5B
PTMA
LOC480914
CDH23
PCCA
JUNB
CD79A
TMSB10
EHD3
MCC
LGALS1
MIF
CXXC5
RPS26
LOC606860
HARBI1
GCNT1
CBLB
CRIP1
TOX
LGALS3
S100A11
ADK
KLF2
ITGB1
TNFRSF4
SYNE2
IL2RA
ZC3H12D
HPCAL1
FAU
LOC475935
RPL28
ZCCHC2
OPTN
LOC100688793
CD4_CTL
LOC483396
IL12RB2
LOC475935
IL18R1
LOC100686073
KLRG1
ZBTB16
ANTXR2
PRKX
MAPKAPK2
ST3GAL6
RPS11
IL7R
TNFAIP3
LOC484314
EPAS1
VIM
LOC102156069
LGALS3
ZFP36L2
SYTL3
PRUNE2
GALNT17
SLC9A9
DOCK3
BHLHE40
MAN1A1
AHNAK
FGL2
IL2RB
CD9
FAR2
BCL2A1
S100A10
CCR5
NKG7
CD99
LCP2
GAPDH
LOC111097825
CD226
RPS2
OSBPL6
ANOS1
CISH
RPL3
DUSP1
JUNB
IL18RAP
GZMK
DN_CTL
ZEB2
RAP1GAP2
DOCK5
SRGAP3
NKG7
GZMB
AOAH
PRKX
SYNE1
FAM53B
KLRD1
PICALM
ATP8A1
PLCG2
YBX3
PREX1
PARP8
PRKCH
ACOXL
CHST11
ARAP2
CLNK
CCL5
CYRIA
RASAL2
PLCB1
SRGAP2
AHNAK
GZMK
SNTB1
PTPRE
JARID2
NCR3
FCER1G
KLRK1
CFD
LOC119869317
ABR
CHN2
TTBK2
DPYD
PSAP
STK39
GZMA
LOC111090626
CD96
ABCA1
TRPS1
ATXN1
CSF1
DN_Tcell
KANK1
NMB
SESN3
PRKCE
AFF3
KIAA0825
GZMK
FOXP1
SYNJ2
PLCL1
BICDL1
TMEM132D
CTSW
TRPS1
KLF12
EPB41L2
IKZF3
WIPF3
ATP8A1
APP
LOC119870193
ICA1
MYB
LOC608120
CBLB
GFOD1
BACH2
LOC606860
SLF1
DTNB
TLK1
ATXN1
KCNQ5
CAMK2D
UBASH3B
PVRIG
DIS3L2
UBE2E2
ADIRF
POU2F2
CASP3
SNX9
VPS13B
RASAL2
ATAD3A
CERS6
ABHD17B
FAF1
FAM102B
MCTP1
gd_Tcell
PSD3
CDH4
PDE11A
RHEX
LOC611565
GATA3
LOC119863905
SYTL3
CRLF2
IL17RB
CMA1
LOC102152421
KCNQ5
BLK
LOC100684807
ELOVL5
P2RY14
LOC111094591
PARD3B
LOC119866429
INPP4B
MYCBP2
GALNT10
MSRB3
ZBTB20
GSTA4
REV3L
XYLT1
RAPGEF5
NFKB1
PDE3B
LOC111095611
PIP5K1B
TNS3
JAZF1
PAG1
C1H9orf3
CSGALNACT1
RIPOR2
SOX13
TMEM273
TRAF3IP2
INPP5D
TCF12
LGALS3
MAML2
CTSW
SNTB1
CDC14A
SCEL
MAIT
IL23R
IL1R1
LOC491694
HDAC9
ZBTB16
PERP
LOC111094591
LOC111093624
PPM1H
ID2
KLRB1
PI3
TNS3
ADAM12
BLK
LOC102155496
DSTN
CLNK
LOC100686073
PTPN13
PLCB1
CPNE8
INPP5D
ANKRD6
RHEX
IKZF2
LOC111095914
AKAP12
NOTCH2
KCTD12
LGALS3
SEPTIN11
SLC9A9
LOC111093623
SNTB1
CDO1
CADM1
GPX1
LOC111092577
ELK3
LOC111090473
RARRES2
ARHGAP32
HMGN3
AAK1
RORC
CCR6
SPOCK1
LOC102156069
CCDC88A
CD4_Th1
CD4_Treg
CD4_CTL
DN_CTL
DN_Tcell
gd_Tcell
MAIT

**Figure S4: Transcriptional signature dynamics of CD4$^{+}$ and DN T cells populations across CanL stages. Related to Figure 4.** Dot plots display gene expression and cell proportions across CanL clinical stages for each CD4$^{+}$/doublet-negative (DN) subset, indicated by their border colors. The dynamics of the top 50 DEGs defining the signature are summarized with line graphs showing the mean Z-score (dark wine-red line) and its standard deviation (light wine-red ribbon). These trajectories are visualized alongside a control distribution (grey ribbon) representing the upper and lower confidence intervals generated by bootstrapping (n = 100,000) non-signature genes.

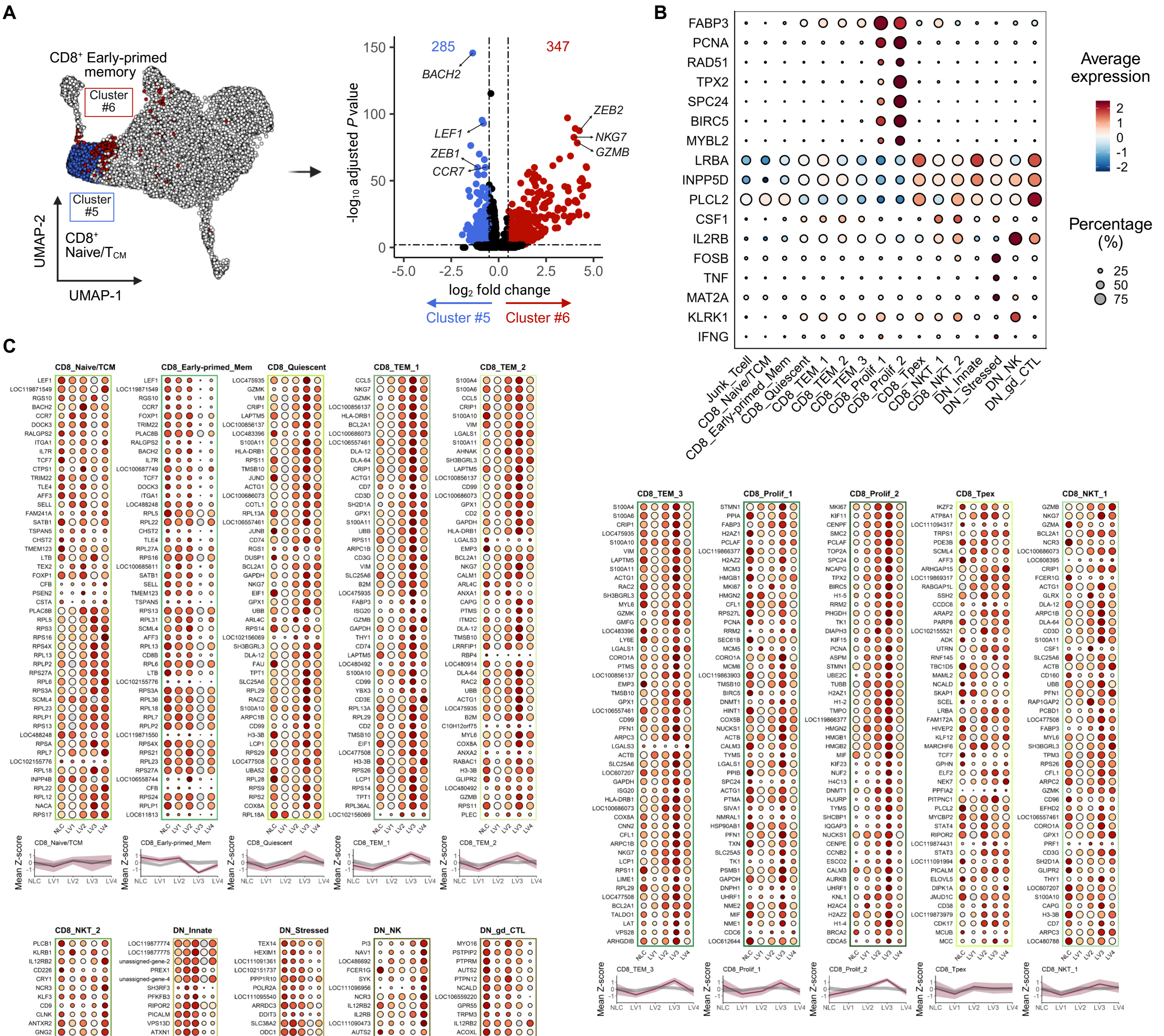

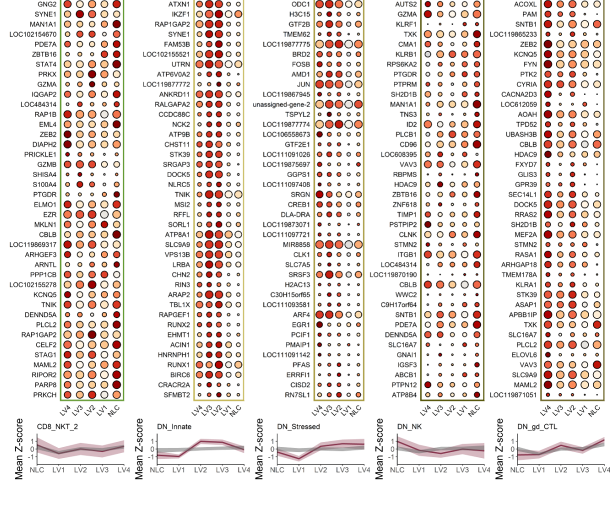

**Figure S5: Differential expression, canonical marker profiles, and transcriptional signature dynamics of CD8⁺ and cytotoxic DN T/NK cell subsets across CanL stages. Related to Figure 5.**

**(A)** Volcano plot showing differential expression between CD8⁺ Early-primed memory (cluster 6) and naïve/TCM (cluster 5) T cell subsets ($|\log_2FC| > 0.5$, adjusted $P < 0.01$).

**(B)** Dot plot showing additional canonical marker expression and proportions of annotated CD8+ and Cyto_DN subclusters.

**(C)** Dot plots display gene expression and cell proportions across CanL clinical stages for each CD8⁺/Cyto_DN subset, indicated by their border colors. The dynamics of the top 50 DEGs defining the signature are summarized with line graphs showing the mean Z-score (dark wine-red line) and its standard deviation (light wine-red ribbon). These trajectories are visualized alongside a control distribution (grey ribbon) representing the upper and lower confidence intervals generated by bootstrapping (n = 100,000) non-signature genes.

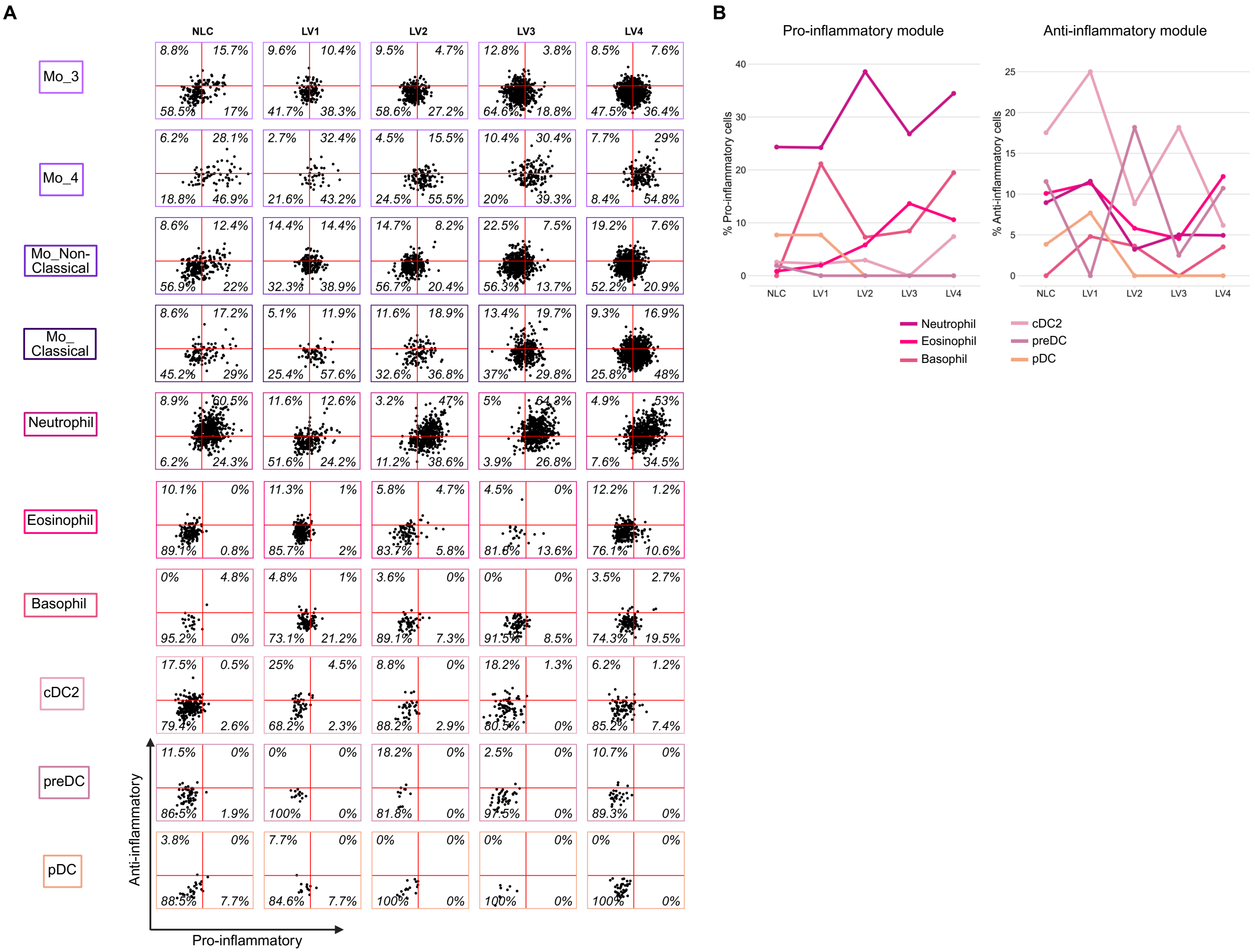

**Figure S6: Inflammatory profiling of myeloid populations. Related to Figure 6.**

**(A)** Scatter plots of inflammatory profiling that juxtapose pro- and anti- inflammatory module scores of Mo_3, Mo_4, Mo_Non-Classical, Mo_Classical, Neutrophil, Eosinophil, Basophil, cDC2, preDC and pDC cell populations at different clinical stages of CanL. Plots classify cells as pro-inflammatory (top left), anti-inflammatory (bottom right) non-inflammatory (bottom left) or mixed inflammatory (top right); and show percentage of cells in each quartile.

**(B)** Line graphs summarizing inflammatory profiling of Neutrophil, Eosinophil, Basophil, cDC2, preDC and pDC populations across CanL clinical stage. The graphs visualize percentages of cells classified as pro- or anti-inflammatory at different clinical stages of CanL.

Table S1. Cohort demographics and bloodwork. Overview of the age, sex, serum chemistry, and blood counts by LeishVet status.

| | Clinical stage | | | | |
|---|---|---|---|---|---|
| | **NLC** | **LV1** | **LV2** | **LV3** | **LV4** |
| | *N* = 3[a] | *N* = 2[b] | *N* = 6 | *N* = 3 | *N* = 2 |
| ***Demographics*** | | | | | |
| *Age (years)* | | | | | |
| Range (%) | | | | | |
| Juvenile (0-2) | 0 | 2 (100) | 1 (16.67) | 1 (33.33) | 0 |
| Adult (3-5) | 2 (75) | 0 | 4 (66.67) | 1 (33.33) | 2 (100) |
| Elderly (≥6) | 0 | 0 | 1 (16.67) | 1 (33.33) | 0 |
| Unknown | 1 (25) | 0 | 0 | 0 | 0 |
| Mean (SD) | 3.5 (0.707) | 1 (0) | 4.167 (1.329) | 4.333 (2.082) | 3.5 (0.707) |
| *Sex (%)* | | | | | |
| Male | 1 (33.33) | 1 (50) | 5 (83.33) | 2 (66.67) | 2 (100) |
| Female | 1 (33.33) | 1 (50) | 1 (16.67) | 1 (33.33) | 0 |
| Unknown | 1 (33.33) | 0 | 0 | 0 | 0 |
| *Tick-serology exposure (%)* | | | | | |
| No | 0 | 0 | 3 (50) | 1 (33.33) | 0 |
| Yes | 0 | 0 | 3 (50) | 2 (66.67) | 2 (100) |
| ***Bloodwork*** | | | | | |
| *Creatinine* | | | | | |
| Mean (SD) (0.5-1.5 mg/dL) | N/A | 1 (0) | 0.633 (0.225) | 1.467 (1.343) | **3.650 (1.061)** |
| *BUN* | | | | | |
| Mean (SD) (9-31 mg/dL) | N/A | 20 (0) | 13.0 (2.280) | **55.67 (69.06)** | **81.0 (26.87)** |
| *Total protein* | | | | | |
| Mean (SD) (5.5-7.5 g/dL) | N/A | 5.9 (0) | **8.733 (1.209)** | **8.967 (0.451)** | **9.85 (2.192)** |
| *Albumin* | | | | | |
| Mean (SD) (2.7-3.9 g/dL) | N/A | 3.2 (0) | **1.617 (0.688)** | **1.633 (0.305)** | **1.45 (0.212)** |
| *Globulin* | | | | | |
| Mean (SD) (2.5-4.0 g/dL) | N/A | 2.7 (0) | **7.117 (1.663)** | **7.333 (0.473)** | **8.4 (2.404)** |
| *ALB/GLOB ratio* | | | | | |
| Mean (SD) (0.7-1.5) | N/A | 1.2 (0) | **0.267 (0.225)** | **0.267 (0.0577)** | **0.15 (0.0707)** |

| | | | | | |
|---|---|---|---|---|---|
| *ALT* | | | | | |
| Mean (SD) (18-121 U/L) | N/A | 22 (0) | 46.17 (30.05) | **17.33 (1.155)** | 24.50 (19.09) |
| *ALP* | | | | | |
| Mean (SD) (5-160 U/L) | N/A | 55 (0) | 56.17 (33.33) | 34.33 (22.37) | 28.50 (6.364) |
| *RBCs* | | | | | |
| Mean (SD) (5.39-8.70 M/µL) | N/A | 6.46 (0) | 6.014 (1.237) | **3.87 (1.023)** | **3.665 (0.46)** |
| *Hematocrit* | | | | | |
| Mean (SD) (38.3-56.5%) | N/A | 45 (0) | **38.07 (10.60)** | **27.97 (7.184)** | **27.10 (1.131)** |
| *Hemoglobin* | | | | | |
| Mean (SD) (13.4-20.7 g/dL) | N/A | 15.80 (0) | 12.88 (3.640) | **8.733 (2.230)** | **8.6 (0.1414)** |
| *WBCs* | | | | | |
| Mean (SD) (4.9-17.6 K/µL) | N/A | 10.7 (0) | **18.9 (3.552)**[c] | 13.53 (6.332) | **18.65 (8.556)** |
| *Retics* | | | | | |
| Mean (SD) (10-110 K/µL) | N/A | 45 (0) | 53.80 (22.24) | 50.67 (17.01) | 24.50 (12.02) |
| *Platelets* | | | | | |
| Mean (SD) (143-448 K/µL) | N/A | 229 (0) | 233.5 (58.38) | 194.3 (76.96) | **133 (0)** |

Bolded values indicate the mean is outside of the normal reference range.
ALB: albumin; ALP: alkaline phosphatase; ALT: alanine transaminase; BUN: blood urea nitrogen; CREA: creatinine; GLOB: globulin; LV: LeishVet score; NLC: non-*Leishmania* control; RBC: red blood cells; Retics: reticulocytes; SD: standard deviation; WBC: white blood cells.
[a]Pooled-sample case considered as missing value (unknown), except for tick-borne exposure.
[b]Only one LV1 dog had a full blood chemistry panel performed.
[c]Complete WBC was available for five of the six LV2 dogs.

Table S2. Frequencies (%) of each PBMC, B, T/NK, CD4$^{+}$ & DN T, CD8$^{+}$ & cytotoxic DN T/NK, and myeloid population across clinical stages. Related to Figures 1-6.

| **Population** | **Clinical stage** | | | | |
|---|---|---|---|---|---|
| | **NLC** | **LV1** | **LV2** | **LV3** | **LV4** |
| ***PBMC (Level 1)*** | | | | | |
| T/NK | 58.41 | 74.41 | 72.23 | 58.00 | 45.32 |
| B | 14.67 | 9.83 | 13.73 | 10.87 | 11.42 |
| Myeloid | 26.92 | 15.76 | 14.05 | 31.13 | 43.26 |
| ***B cells (Level 2)*** | | | | | |
| Transitional | 4.13 | 9.72 | 6.28 | 4.54 | 3.39 |
| Naive | 51.38 | 59.67 | 60.75 | 54.12 | 59.36 |
| Unswitched | 14.03 | 11.08 | 10.56 | 11.04 | 9.99 |
| B_1 | 20.65 | 9.20 | 11.94 | 22.22 | 17.39 |
| B_2 | 6.11 | 4.81 | 7.79 | 5.90 | 5.91 |
| Plasma cells | 3.70 | 5.54 | 2.68 | 2.19 | 3.96 |
| ***T/NK cells (Level 2)*** | | | | | |
| CD4$^{+}$ | 64.82 | 56.91 | 34.20 | 34.50 | 44.75 |
| CD8$^{+}$ | 20.50 | 26.38 | 50.54 | 49.33 | 38.87 |
| DN | 9.00 | 8.32 | 6.34 | 7.81 | 7.08 |
| Cytotoxic DN | 5.68 | 8.39 | 8.92 | 8.36 | 9.29 |
| ***CD4$^{+}$ & DN T cells (Level 3)*** | | | | | |
| CD4_Naive | 32.82 | 45.55 | 29.44 | 20.12 | 24.66 |
| CD4_TCM | 5.28 | 7.26 | 4.78 | 5.91 | 4.85 |
| CD4_IFN_stim | 1.83 | 2.86 | 2.68 | 3.43 | 2.21 |
| CD4_Int | 13.08 | 10.55 | 8.98 | 11.87 | 12.72 |
| CD4_$T_{EM}$ | 14.08 | 8.01 | 11.67 | 12.58 | 10.51 |
| CD4_$T_{H}$2 | 8.28 | 5.38 | 9.16 | 15.53 | 12.07 |
| CD4_$T_{H}$17 | 3.83 | 3.03 | 3.33 | 6.64 | 8.03 |
| CD4_$T_{H}$1 | 1.17 | 1.17 | 2.45 | 4.14 | 3.83 |
| CD4_$T_{REG}$ | 3.39 | 1.80 | 2.14 | 2.79 | 2.43 |
| CD4_CTL | 5.11 | 3.62 | 3.87 | 4.49 | 3.85 |
| DN_CTL | 4.78 | 2.25 | 14.16 | 7.86 | 11.40 |
| DN_Tcell | 1.78 | 1.12 | 2.53 | 1.64 | 2.21 |
| gd_Tcell | 4.28 | 6.95 | 4.24 | 2.54 | 1.19 |
| MAIT | 0.31 | 0.45 | 0.58 | 0.45 | 0.05 |
| ***CD8$^{+}$ & cytotoxic DN T/NK cells (Level 3)*** | | | | | |
| CD8_Naive/$T_{CM}$ | 10.58 | 10.78 | 5.00 | 4.17 | 7.20 |
| CD8_Early-primed_Mem | 13.58 | 9.84 | 5.21 | 1.60 | 3.54 |
| CD8_Quiescent | 18.75 | 15.44 | 15.01 | 12.76 | 15.21 |

| | | | | | |
|---|---|---|---|---|---|
| CD8_TEM_1 | 9.42 | 13.83 | 19.82 | 22.14 | 19.68 |
| CD8_TEM_2 | 2.42 | 4.11 | 5.99 | 5.65 | 4.91 |
| CD8_TEM_3 | 6.33 | 4.94 | 3.68 | 4.70 | 3.75 |
| CD8_Prolif_1 | 0.75 | 1.37 | 2.10 | 3.46 | 2.29 |
| CD8_Prolif_2 | 2.42 | 1.25 | 1.27 | 2.29 | 2.14 |
| CD8_$T_{PEX}$ | 9.42 | 7.76 | 11.00 | 10.64 | 8.43 |
| CD8_NKT_1 | 6.67 | 7.64 | 9.12 | 8.87 | 9.47 |
| CD8_NKT_2 | 5.25 | 5.49 | 8.73 | 10.33 | 6.01 |
| DN_Innate | 1.50 | 2.74 | 5.49 | 5.53 | 3.39 |
| DN_Stressed | 4.08 | 3.13 | 4.85 | 3.53 | 4.76 |
| DN_NK | 4.58 | 3.57 | 0.99 | 1.74 | 0.60 |
| DN_gd_CTL | 4.25 | 8.11 | 1.74 | 2.58 | 8.60 |
| ***Myeloid cells (Level 2)*** | | | | | |
| Mo_1 | 13.55 | 13.48 | 18.18 | 17.39 | 21.55 |
| Mo_2 | 12.27 | 15.73 | 19.96 | 21.26 | 18.49 |
| Mo_3 | 8.13 | 8.61 | 11.16 | 13.27 | 14.00 |
| Mo_4 | 3.27 | 2.77 | 5.29 | 3.96 | 2.33 |
| Mo_Non-Classical | 10.69 | 12.51 | 11.78 | 15.27 | 12.56 |
| Mo_Classical | 4.75 | 4.42 | 4.57 | 8.96 | 14.89 |
| Neutrophil | 26.28 | 14.23 | 19.34 | 13.48 | 8.49 |
| Eosinophil | 6.08 | 15.21 | 4.14 | 0.65 | 3.83 |
| Basophil | 1.07 | 7.79 | 2.65 | 2.09 | 1.70 |
| cDC2 | 9.92 | 3.30 | 1.64 | 2.26 | 1.22 |
| preDC | 2.66 | 0.97 | 0.53 | 1.17 | 0.42 |
| pDC | 1.33 | 0.97 | 0.77 | 0.23 | 0.53 |

cDC: conventional dendritic cell; CM: central memory; CTL: cytotoxic T lymphocyte; DN: double-negative; EM: effector memory; gd: gamma-delta; pDC: plasmacytoid dendritic cell; preDC: precursor dendritic cell; NLC: non-*Leishmania* control; MAIT: mucosal associated-invariant T; Mo: monocyte; NK: Natural Killer; PBMC: peripheral blood mononuclear cells; PEX: progenitor exhaustion.

Table S3. Fold-changes in log odds ratios comparing abundances of peripheral immune cell types and major T/NK cell subsets across CanL stages versus NLC. Related to Figures 1 and 3.

| Contrast | LV score | *Z* ratio | FC OR | CI | Adjusted *P* |
|---|---|---|---|---|---|
| ***PBMC (Level 1)*** | | | | | |
| T/NK : Myeloid | LV1 | 29.69 | 4.0760844 | [3.72, 4.47] | 1.26e-193 |
| T/NK : Myeloid | LV2 | 34.00 | 4.1735447 | [3.84, 4.53] | 9.54e-253 |
| T/NK : Myeloid | LV3 | -5.33 | 0.801078 | [0.74, 0.87] | 9.98e-08 |
| T/NK : Myeloid | LV4 | -31.90 | 0.2850927 | [0.26, 0.31] | 6.20e-223 |
| T/NK : B | LV1 | 21.90 | 3.2650699 | [2.94, 3.63] | 1.07e-105 |
| T/NK : B | LV2 | 15.11 | 2.0019679 | [1.83, 2.19] | 2.84e-51 |
| T/NK : B | LV3 | 6.54 | 1.3867489 | [1.26, 1.53] | 8.32e-11 |
| T/NK : B | LV4 | -5.07 | 0.7871008 | [0.72, 0.86] | 4.00e-07 |
| Myeloid : B | LV1 | -3.90 | 0.801031 | [0.72, 0.90] | 9.74e-05 |
| Myeloid : B | LV2 | -15.01 | 0.479681 | [0.44, 0.53] | 1.36e-50 |
| Myeloid : B | LV3 | 10.65 | 1.7311035 | [1.56, 1.92] | 2.36e-26 |
| Myeloid : B | LV4 | 20.97 | 2.7608586 | [2.51, 3.04] | 4.63e-97 |
| ***T/NK cells (Level 2)*** | | | | | |
| $CD4^+$ : $CD8^+$ | LV1 | -11.46 | 0.51577560 | [0.46, 0.58] | 1.98e-30 |
| $CD4^+$ : $CD8^+$ | LV2 | -49.99 | 0.07121129 | [0.06, 0.08] | 0.00e+00 |
| $CD4^+$ : $CD8^+$ | LV3 | -44.92 | 0.07572185 | [0.07, 0.08] | 0.00e+00 |
| $CD4^+$ : $CD8^+$ | LV4 | -29.98 | 0.17826158 | [0.16, 0.20] | 2.70e-197 |
| $CD4^+$ : DN | LV1 | -3.31 | 0.7805136 | [0.67, 0.90] | 0.0009 |
| $CD4^+$ : DN | LV2 | -12.52 | 0.4123549 | [0.36, 0.47] | 1.16e-35 |
| $CD4^+$ : DN | LV3 | -14.30 | 0.3338367 | [0.29, 0.39] | 9.29e-46 |
| $CD4^+$ : DN | LV4 | -7.19 | 0.5704599 | [0.49, 0.66] | 8.46e-13 |
| $CD4^+$ : Cytotoxic DN | LV1 | -9.05 | 0.4712033 | [0.40, 0.55] | 1.39e-19 |
| $CD4^+$ : Cytotoxic DN | LV2 | -22.64 | 0.1733844 | [0.15, 0.20] | 6.57e-113 |
| $CD4^+$ : Cytotoxic DN | LV3 | -19.80 | 0.1885554 | [0.16, 0.22] | 5.79e-87 |
| $CD4^+$ : Cytotoxic DN | LV4 | -16.23 | 0.2584756 | [0.22, 0.30] | 4.13e-59 |
| $CD8^+$ : DN | LV1 | 5.31 | 1.513281 | [1.30, 1.76] | 1.11e-07 |
| $CD8^+$ : DN | LV2 | 24.05 | 5.790584 | [5.02, 6.68] | 3.25e-127 |
| $CD8^+$ : DN | LV3 | 18.86 | 4.408724 | [3.78, 5.14] | 5.27e-79 |
| $CD8^+$ : DN | LV4 | 14.46 | 3.200128 | [2.73, 3.75] | 2.92e-47 |

| | | | | | |
|---|---|---|---|---|---|
| CD8$^+$ : Cytotoxic DN | LV1 | -1.05 | 0.913582 | [0.77, 1.08] | 0.2935 |
| CD8$^+$ : Cytotoxic DN | LV2 | 11.20 | 2.434788 | [2.08, 2.85] | 1.66e-28 |
| CD8$^+$ : Cytotoxic DN | LV3 | 10.61 | 2.490106 | [2.10, 2.95] | 5.62e-26 |
| CD8$^+$ : Cytotoxic DN | LV4 | 4.34 | 1.449980 | [1.23, 1.71] | 1.90e-05 |
| DN : Cytotoxic DN | LV1 | -5.13 | 0.6037093 | [0.50, 0.73] | 2.85e-07 |
| DN : Cytotoxic DN | LV2 | -9.38 | 0.4204737 | [0.35, 0.50] | 2.59e-20 |
| DN : Cytotoxic DN | LV3 | -5.72 | 0.5648133 | [0.46, 0.69] | 1.46e-08 |
| DN : Cytotoxic DN | LV4 | -7.87 | 0.4531005 | [0.37, 0.55] | 6.94e-15 |

DN: double-negative; LV: LeishVet; NLC: non-*Leishmania* control; PBMC: peripheral blood mononuclear cells.
